# Combinatorial gene inactivation of aldehyde dehydrogenases mitigates aldehyde oxidation catalyzed by resting cells of *E. coli* RARE strains

**DOI:** 10.1101/2023.01.16.524286

**Authors:** Neil D. Butler, Shelby R. Anderson, Roman M. Dickey, Priyanka Nain, Aditya M. Kunjapur

## Abstract

Aldehydes are attractive chemical targets given applications as end products in the flavors and fragrances industry and as intermediates due to their propensity for C-C bond formation. While biosynthetic routes to diverse aldehydes have been designed, a common challenge is the stability of these aldehydes in the presence of microbial hosts of engineered pathways. Here, we identify and address unexpected oxidation of a model collection of aromatic aldehydes, including many that originate from biomass degradation, in the presence of *Escherichia coli* strains that were engineered to minimize aldehyde reduction. Of heightened interest to us were resting cell conditions as they offer numerous advantages for the bioconversion of toxic metabolites. Surprisingly, when diverse aldehydes are supplemented to *E. coli* RARE cells grown under aerobic conditions, they remain stabilized on the timescale of days, whereas when these same aldehydes are supplemented to resting cell preparations of *E. coli* RARE that had been grown under the same conditions, we observe substantial oxidation. By performing combinatorial inactivation of six candidate aldehyde dehydrogenase genes in the *E. coli* genome using multiplexed automatable genome engineering (MAGE), we demonstrate that this oxidation can be substantially slowed, with greater than 50% retention of 6 out of 8 aldehydes when assayed 4 hours after their addition. Given that our newly engineered strain exhibits <u>R</u>educed <u>O</u>xidation <u>A</u>nd <u>R</u>eduction of aromatic aldehydes, we dubbed it the *E. coli* ROAR strain. Seeking to apply this new strain to resting cell biocatalysis, we compared the capability to synthesis the aldehyde furfural from 2-furoic acid via the carboxylic acid reductase enzyme from *Nocardia iowensis*. Here, we found that use of ROAR resting cells achieved 2-fold enhancement in furfural titer after 4 h and nearly 9-fold enhancement after 20 h as compared to resting cells of the RARE strain. Moving forward, the use of this strain to generate resting cells should allow aldehyde product isolation, further enzymatic conversion, or chemical reactivity under cellular contexts that better accommodate aldehyde toxicity.

**Highlights:**

- When genes that encode aldehyde reductases are knocked out in *Escherichia coli* strains, supplemented aldehydes can experience oxidation instead of reduction, which is catalyzed by a different set of endogenous enzymes.
- Interestingly, we show for a collection of aromatic aldehydes that this oxidation is far more substantial when using resting cell preparations than during aerobic fermentation.
- We investigate the identity of the responsible genes by performing combinatorial gene inactivation using multiplex automatable genome engineering.
- The strain that we engineer exhibits Reduced Oxidation And Reduction (the *E. coli* ROAR strain) and thereby enables design of more efficient aldehyde bioconversion processes under diverse formats.

## 1. Introduction

Over the last fifteen years, the biosynthesis of aldehydes has risen to prominence across diverse scientific disciplines. The biosynthesis of aldehydes as end products is of high active interest in industry (Kunjapur and Prather, 2015), and notable examples reported in the academic literature include the biosynthesis of vanillin from simple carbon sources (Hansen et al., 2009; Kunjapur et al., 2014) or from supplemented precursors (Gallage et al., 2014; Sadler and Wallace, 2021) as well as biosynthesis of many other aldehyde food and fragrance compounds such as benzaldehyde (Duff and Murray, 1989; Kunjapur et al., 2016, 2014; Orbegozo et al., 2009), piperonal (Schwendenwein et al., 2016), octanal (Horvat and Winkler, 2020), and cinnamaldehyde (Son et al., 2022) from supplemented precursors using microbial cells or enzymes. However, the last decade has given rise to a plethora of new opportunities for aldehydes to serve as intermediates in biosynthetic pathways or enzyme cascades. Aliphatic aldehydes generated from fermentation or fatty acid synthesis have been used to produce fuels and commodities based on alcohols (Akhtar et al., 2013; Sheppard et al., 2014) or alkanes (Kallio et al., 2014; Sheppard et al., 2016). Exemplary newer aldehyde-derived biosynthetic targets that have applications in the pharmaceutical and materials industries are tetrahydroisoquinoline alkaloid plant natural products (Pyne et al., 2020), primary or heterocyclic mono-amine precursors to small molecule pharmaceuticals (Citoler et al., 2019; France et al., 2016; Hepworth et al., 2017), diamine polymer building blocks (Fedorchuk et al., 2020; Gopal et al., 2022), hydroxylated non-standard amino acids (Doyon et al., 2022; Ellis et al., 2022; Kumar et al., 2021; Xu et al., 2020, 2019), and β-lactone antibiotics (Schaffer et al., 2017; Scott et al., 2017). Additionally, opportunities to blend biological and abiological synthesis methods to access additional chemical functional groups are emerging, such as the recent chemoenzymatic synthesis of nitriles (Horvat et al., 2022; Winkler et al., 2022).

As these examples illustrate, aldehydes have served as a gateway to C-C bond formation and heteroatom introduction due to the electrophilicity of the carbonyl group, enabling diversification of structure and chemical functional group (Hanefeld et al., 2022; Sulzbach and Kunjapur, 2020). In addition to the biosynthesis of small molecule aldehydes, the biosynthesis of aldehydes on the residues of proteins has also been of high academic and commercial relevance since the demonstration by Bertozzi and colleagues of introducing a genetically encoded aldehyde tag within protein sequences (Carrico et al., 2007). The post-translational formation of formylglycine residues within proteins enable their biologically orthogonal conjugation to small molecules to create conjugate protein therapies. Efforts to improve the cellular environment for aldehyde biosynthesis may offer benefits for protein applications that feature aldehydes.

While certain aldehydes are present in natural fermentation pathways and in secondary metabolism, the relevance of aldehydes to metabolic engineers has increased in recent years in large part due to the identification of carboxylic acid reductases (CARs). CARs are large (∼130 kDa), multi-domain enzymes which can perform two electron reductions of a broad range of both aromatic and aliphatic carboxylic acid substrates (Gahloth et al., 2020). The polyspecificity and generally high activity of CARs have enabled the conversion of numerous cellular metabolites or supplemented precursors to their aldehyde form by bacterial cells after heterologous expression (Butler and Kunjapur, 2020). However, accomplishing effective biocatalysis with CARs necessarily requires co-expression of a separate enzyme for post-translational modification (phosphopantetheine transferase, or PPTase) and sufficient pools of the cofactors ATP and NADPH. Given these constraints, researchers have pursued several different avenues to optimize CAR activity depending on the enzyme’s native expression or activity on a substrate of interest such as *in vitro* cofactor recycling cascades, resting whole cell biocatalysis, or metabolic engineering to maximize cofactor availability and small molecule aldehyde production *in vivo*.

Despite the substantial advances in the ability to biosynthesize aldehydes, two important challenges that limit the ability to feature small molecule aldehydes in cellular environments are their instability due to the activity of cellular enzymes and their toxicity to microbial cells (Dickey et al., 2021). In the last decade, the deletion of some of the many native genes that encode aldehyde reductases in *E. coli* resulted in dramatic improvements in stability for several aromatic and aliphatic aldehydes under aerobic culturing conditions (Kunjapur et al., 2014; Rodriguez and Atsumi, 2014). The deletion of multiple such genes in yeast also contributed to significant improvements in aldehyde stability as observed indirectly through (*S*)-norcoclaurine biosynthesis (Pyne et al., 2020). While at first glance these gene deletion studies appear to have mostly remedied the issue of aldehyde stability for several aldehydes under certain conditions, we sought to test this assumption in more detail. We were particularly curious about aldehyde stability under conditions that could be more relevant for addressing the outstanding issue of aldehyde toxicity. While aldehyde intermediates may be kept at a low steady-state concentration due to the action of downstream enzymes, aldehyde products interfere with cell growth if not immediately separated.

One compelling alternative to the concept of producing aldehyde end products during fermentative growth is to instead harness resting whole cell biocatalysts. The use of such resting cells shares similarity to the often-employed metabolic engineering strategy of designing a fermentation that features a biomass accumulation phase followed by a production phase. Resting cell bioconversions offer some advantages over fermentative processes, particularly when a precursor cannot be generated from simple carbon sources, as is the case for several aldehydes of interest, or when the toxicity of a product or intermediate would curtail growth (Lin and Tao, 2017). The use of resting cells can also simplify downstream separations as there are fewer fermentative byproducts. Finally, resting cells also offer some advantages over other alternatives such as cell-free enzyme cascades in that they obviate the need for cell lysis, supplementation of expensive co-factors, or laborious protein purification. Notably, resting cells can still be partially metabolically active, as in many cases glucose is provided to cells for regeneration of co-factors such as ATP and NADPH.

In this study, we investigated the stability of biomass-derived aromatic aldehydes when supplemented to either *E. coli* K-12 MG1655 or the engineered RARE (**r**educed aromatic **a**ldehyde **re**duction) strain in the absence of CAR expression to reveal potential instability in the oxidative direction. We examined conditions of aerobic fermentative growth and resting cells prepared after aerobic fermentative growth. Using the RARE strain, we reveal significant differences in the tendency of cells to catalyze aldehyde oxidation under these conditions. We also use multiplex automatable genome engineering (MAGE) to perform translational knockouts of several aldehyde dehydrogenases that may contribute to this activity. Our study achieved significant decreases in the rate of aldehyde oxidation, which we then showed can lead to large improvements in aldehyde titer arising from resting cell bioconversions.

## 2. Materials and Methods

### 2.1. Strains and plasmids

*Escherichia coli* strains and plasmids used are listed in **Table S1**. Molecular cloning and vector propagation were performed in DH5α. Polymerase chain reaction (PCR) based DNA replication was performed using KOD XTREME Hot Start Polymerase for plasmid backbones or using KOD Hot Start Polymerase otherwise. Cloning was performed using Gibson Assembly. Oligos for PCR amplification and translational knockouts are shown in **Table S2**. Oligos were purchased from Integrated DNA Technologies (IDT). The pORTMAGE-EC1 recombineering plasmid was kindly provided by Timothy Wannier and George Church of Harvard Medical School.

### 2.2. Chemicals

The following compounds were purchased from MilliporeSigma: kanamycin sulfate, dimethyl sulfoxide (DMSO), imidazole, potassium phosphate dibasic, potassium phosphate monobasic, magnesium sulfate, anhydrous magnesium chloride, calcium chloride dihydrate, glycerol, M9 salts, lithium hydroxide, boric acid, Tris base, glycine, HEPES, ATP, 2-napthaldehyde, 4-anisaldehyde, 3-hydroxybenzaldehyde, syringaldehyde, vanillin, benzaldehyde, furfural, 4-hydroxy-3-methoxybenzyl alcohol, furfuryl alcohol, 4-methoxybenzoic acid, 3-hydroxybenzoic acid, syringic acid, vanillic acid, benzoic acid and KOD XTREME Hot Start and KOD Hot Start polymerases. D-glucose, m-toluic acid, piperonal, piperonyl alcohol, 2-naphtol, 4-methoxybenzyl alcohol, 3-hydroxybenzyl alcohol, piperonylic acid, 2-naphtoic acid, and 2-furoic acid were purchased from TCI America. Agarose and ethanol were purchased from Alfa Aesar. Acetonitrile, sodium chloride, LB Broth powder (Lennox), LB Agar powder (Lennox), were purchased from Fisher Chemical. A MOPS EZ rich defined medium kit was purchased from Teknova. Trace Elements A was purchased from Corning. Taq DNA ligase was purchased from GoldBio. Phusion DNA polymerase and T5 exonuclease were purchased from New England BioLabs (NEB). Sybr Safe DNA gel stain was purchased from Invitrogen. Benzyl alcohol was purchased from Merck. 4-hydroxy-3,5-dimethoxybenzoic alcohol was purchased from ChemCruz. NADPH (tetrasodium salt) was purchased from Santa Cruz Biotechnology. Anhydrotetracycline was purchased from Cayman Chemical.

### 2.3. Culture Conditions

Cultures were grown in LB-Lennox medium (LBL: 10 g/L bacto tryptone, 5 g/L sodium chloride, 5 g/L yeast extract), M9-glucose minimal media (Kunjapur et al., 2016) with Corning® Trace Elements A (1.60 µg/mL CuSO_4_ • 5H_2_O, 863.00 µg/mL ZnSO_4_ • 7H_2_O, 17.30 µg/mL Selenite 2Na, 1155.10 µg/mL ferric citrate) and 1.5% glucose, or MOPS EZ rich defined media (Teknova M2105) with 1.5% glucose (MOPS EZ Rich-glucose).

To prepare cells for resting cell assays or cell-free lysate testing, confluent overnight cultures of *E. coli* strains were used to inoculate 100 mL cultures in 500 mL baffled shake flasks. The cultures were grown at 37 °C until mid-exponential phase (OD_600_ = 0.5-0.8), 30 °C for 5 additional hours, and then 18 °C overnight for 18h. Cells were then pelleted and frozen at -80 °C. For growth rate testing, cultures were prepared from confluent overnight cultures used to inoculate experimental cultures at 100x dilution in 200 µL volumes in a Greiner clear bottom 96 well plate (Greiner 655090). Cultures were grown for 20 h in a Spectramax i3x plate reader with medium plate shaking at 37 ºC with absorbance readings at 600 nm taken every 11 min to calculate doubling time and growth rate. For experiments investigating protein expression, strains transformed with a pZE-Ub-sfGFP reporter plasmid were grown as described above with 50 μg/mL Kan and 0.2 nM anhydrotetracycline (aTc) added.

### 2.4. Stability Assays

For metabolically active cell stability testing, cultures of each *E. coli* strain to be tested were inoculated from a frozen stock and grown to confluence overnight in 5 mL of LBL media. Unless otherwise indicated, confluent overnight cultures were then used to inoculate experimental cultures in 400 µL volumes in a 96-deep-well plate (Thermo Scientific™ 260251) at 100x dilution. Cultures were supplemented with 1 mM of heterologous aldehydes (prepared in 100 mM stocks in DMSO) at mid-exponential phase (OD_600_ = 0.5-0.8). Cultures were incubated at 37 °C with shaking at 1000 RPM and an orbital radius of 3 mm. Samples were taken by pipetting 150 µL from the cultures, centrifuging in a round bottom plate (SPL Life Sciences ISO 13485) and collecting the extracellular broth. Compounds were quantified over a 20 h period using HPLC with samples collected at 4 h and 20 h.

For metabolically active cell stability testing in culture tubes, 5 mL of media were inoculated at 100x dilution with confluent overnights in 14 mL culture tubes. Cultures were supplemented with 1 mM of benzaldehyde (prepared in a 100 mM stock in DMSO) at mid-exponential phase (OD∼0.5). Cultures were then incubated at 37 °C in a rotor drum (Thermo Scientific Cel-Gro Tissue Culture Rotator) at maximum speed. Samples were taken via centrifugation after 20 h with compound quantification performed using HPLC.

For resting cell stability testing, cell pellets were thawed and then washed 2x with 200 mM HEPES, pH 7.0 buffer. The mass of cell pellets was then measured, and the pellets were resuspended in 200 mM HEPES, pH 7.0 at a wet cell weight of 50 mg/mL The resuspended resting cells were then aliquoted into 96-deep-well plates at 400 µL and aldehydes of interest (prepared in 100 mM stocks in DMSO) were added at a concentration of 1 mM. Resting cells were then incubated at 37 °C with shaking at 1000 RPM and an orbital radius of 3 mm. Samples were taken by pipetting 200 µL from the cultures, centrifuging in a round bottom plate, and collecting the extracellular broth. Compounds were quantified over a 20 h period using HPLC with samples collected at 4 h and 20 h.

For lysate stability testing, cell pellets were thawed and then washed 2x with 200 mM HEPES, pH 7.0 buffer. Cell pellets were then resuspended in 5 mL of 200 mM HEPES, pH 7.0 and disrupted via sonication using a QSonica Q125 sonicator with cycles of 5 s at 75% amplitude and 10 s off for 5 minutes. The lysate was distributed into microcentrifuge tubes and centrifuged for 1 h at 18,213 x *g* at 4 °C. Supernatant was then collected, and the total protein concentration was then measured using a Bradford assay. Then, the cell-free lysate was concentrated to a final concentration of 5 mg/mL using a 3 kDa cutoff centrifugal concentrator (Amicon Ultra, MilliporeSigma) and aliquoted into 400 µL in 96-deep-well plates and mixed with 1 mM of aldehydes of interest (prepared in 100 mM stocks in DMSO). Cell free lysates were then incubated at 37 °C with shaking at 1000 RPM and an orbital radius of 3 mm. Samples were taken by pipetting 200 µL from the cultures, mixing the sample with 2 µL of 10% trifluoroacetic acid and centrifuging in a round bottom plate, and collecting the extracellular broth. Compounds were quantified after 4 h using HPLC.

### 2.5. Translational genomic knockouts

Translational knockouts to the strain were performed using 10 rounds of multiplexed automatable genome engineering (MAGE) where two stop codons (one TAA and one TGA) were introduced into the genomic sequence for each aldehyde dehydrogenase target in the upstream portion of the gene (within the first 22 amino acid positions). This was performed using the pORTMAGE-EC1 recombineering plasmid (Wannier et al., 2020). Briefly, bacterial cultures were inoculated with 1:100 dilution in 3 mL of LBL media with 50 μg/mL kanamycin (Kan) and grown at 37 °C until OD of 0.4-0.6 was reached. Then, the pORTMAGE plasmid was induced with 1 Mm m-toluic acid and cultures were grown at 37 °C for an additional 10 min. Cells were then prepared for electroporation by washing 1 mL three times with refrigerated 10% glycerol and then resuspending in 50 µL of 10% glycerol with each knockout oligo added at 1 µM. Cells were then electroporated and recovered in 3 mL of LBL with Kan to be used for subsequent rounds. Preliminary assessment of knockouts was performed using PCR and confirmed using Sanger Sequencing. Curing of the pORTMAGE-EC1 plasmid following confirmation of genomic knockouts produced the ROAR strain.

### 2.6. *In vitro* carboxylic acid reductase activity

For purification, the pZE-niCAR-sfp plasmids was transformed into *E. coli* BL21(DE3). 250mL of LB in 1L baffled flasks was supplemented with 50 µg/mL kanamycin and was inoculated 1:100 from confluent overnight culture and grown at 37 °C. At mid-exponential phase (OD_600_ = 0.5-0.8), 0.1 µg/mL aTc was added to induce cultures. Cultures were then grown at 30 °C for 5 h and then 18 °C for 18h. Cells were then harvested by centrifugation.

Next, cells were resuspended in Ni-binding buffer (100 mM HEPES pH 7.4, 250 mM NaCl, 10 mM imidazole, 10% glycerol, 10 mM magnesium chloride). The cells were lysed by sonication followed by centrifugation at 48,000 × g for 1 h. The cleared supernatant was sterile filtered through a 0.22 µm syringe filter and was purified using AKTA Pure fast protein liquid chromatography (FPLC) system containing a Ni-Sepharose affinity chromatography (HisTrap HP, 5mL) with isocratic binding, wash and elution steps. Final elution performed at 250 mM imidazole. Purified fractions were pooled, concentrated, and dialyzed against dialysis buffer (100mM HEPES pH 7.5, 300 mM sodium chloride, 10% glycerol and 10mM magnesium chloride hexahydrate) with 30 kDa molecular weight cutoff centrifugal filters (Amicon Ultra, MilliporeSigma).

The i*n vitro* CAR assay was performed in a buffer containing 200 mM HEPES, pH 7.0 at 30°C, 0.5 mM NADPH, 1 mM ATP, 10 mM MgCl_2_, and 1.5 µM niCAR. To start, 95 µL of the buffer was loaded into a 96 well plate. Prior to spectrophotometric analysis of NADPH depletion, 5 µL of substrate (100 mM in DMSO) was added to each reaction well, resulting in final composition of 5 mM furoic acid and 5% v/v DMSO in 100 µL of reaction mix. Oxidation of NADPH was measured at 30 ºC for triplicate samples using an Agilent BioTek Synergy H4 Hybrid Microplate Reader at a wavelength of 340 nm for 15 minutes. Spectral scans of furfural and furoic acid at 5 mM determined that there was no competing absorbance measurement at 340 nm. *In vitro* CAR activity was then analyzed to determine the maximum rate of NADPH oxidation, using a 0 mM and 0.5 mM NADPH standards as a calibration curve.

### 2.7. Resting cell carboxylic acid reductase screening

To assay niCAR in resting cells, we mixed resting cells concentrated to 50 mg/mL wet cell weight in buffer with 200 mM HEPES, pH 7.0, glucose (either 10 or 100 mM concentration for regeneration of ATP and NADPH), 10 mM magnesium chloride, and 5 mM 2-furoic acid in 1 mL volume in 96-deep well plates. Samples were taken by pipetting 200 µL from the cultures, centrifuging in a round bottom plate, and collecting the extracellular broth. Compounds were quantified over a 20 h period using HPLC.

### 2.8. HPLC analysis

Metabolites of interest were quantified via high-performance liquid chromatography (HPLC) using an Agilent 1100 Infinity model equipped with a Zorbax Eclipse Plus-C18 column (part number: 959701-902, 5 µm, 95Å, 2.1 × 150 mm). To quantify compounds of interest, an initial mobile phase of solvent A/B = 95/5 was used (solvent A, water, 0.1% trifluoroacetic acid; solvent B, acetonitrile, 0.1% trifluoroacetic acid) and maintained for 5 min. A gradient elution was performed (A/B) with: gradient from 95/5 to 50/50 for 5-12 min, gradient from 50/50 to 0/100 for 12-13 min, gradient from 0/100 to 95/5 for 13-14, and equilibration at 95/5 for 14-15 min. A flow rate of 1 mL min^-1^ was maintained, and absorption was monitored at 210, 250, 270, 280 and 300 nm.

## 3. Results

### 3.1. Aldehydes are subject to unexpected and rapid oxidation under certain culturing conditions of E. coli RARE

We first sought to measure the stability of a diverse collection of aromatic aldehydes (**Fig. 1A**) under aerobic growth conditions in a defined medium (**Fig. 1B**). In our previous work, we had biosynthesized aldehydes of interest through heterologous expression of the CAR from *Nocardia iowensis* (niCAR) because of the lower toxicity, lower volatility, increased solubility, and possibly superior uptake of carboxylic acid precursors. However, CAR co-expression could mask potential oxidation catalyzed by native cells. To evaluate whether oxidation occurred in fermentative cultures, we supplemented low (1 mM) concentrations of eight different aldehydes to both *E. coli* K-12 MG1655 and the engineered RARE strain in MOPS EZ Rich-glucose media at mid-exponential phase of growth. To screen these eight aldehydes efficiently, we grew cells of either strain in triplicate, 400 µL cultures in deep 96-well plates sealed with non-breathable aluminum seals to limit loss due to aldehyde volatility. We sampled culture broth at two time points (4 h and 20 h after aldehyde addition) to gain some insight on the kinetics of aldehyde stability and to explore whether it could be a function of growth phase. Under these conditions, we observed mostly expected results. In cultures of the wild-type *E. coli* K-12 MG1655 strain, the supplemented aldehydes were nearly completely reduced to their corresponding alcohol within 4 h (**Fig. 1C**) with this condition persisting after 20 h (**Fig. 1D**).

**Figure 1.**
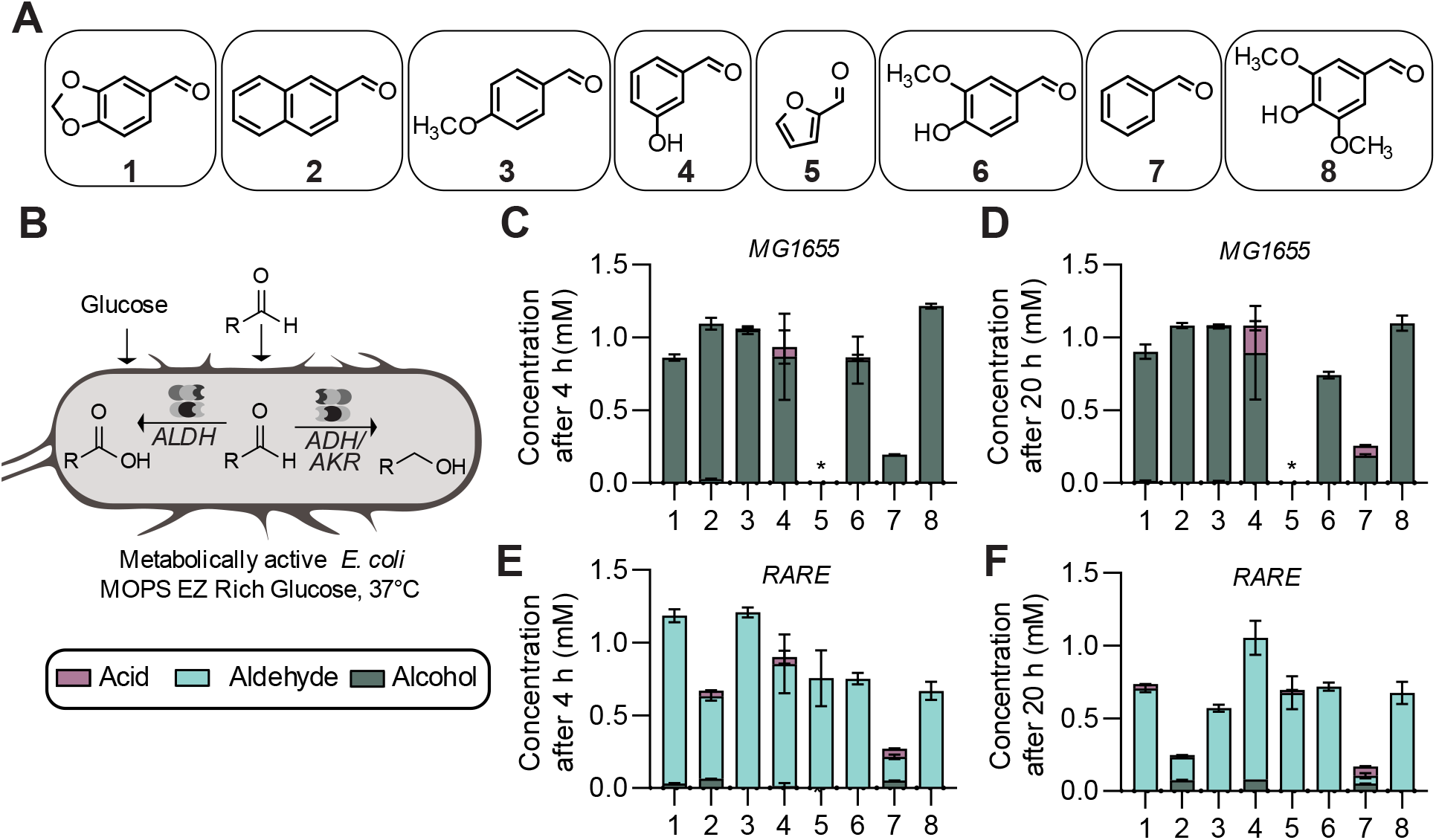
Evaluation of the stability of aldehydes in metabolically active strains. **A**. A panel of aldehydes was selected for their **B**. oxidoreductive stability in metabolically active cultures. **C**. Cultures of wild-type *E. coli* MG1655 were grown in MOPS EZ Rich-glucose media at 37°C with supplementation of 1 mM aldehydes and fate of the aldehydes were tracked via HPLC after 4 h and **D**. 20 h. **E**. Cultures of the engineered *E. coli* strain RARE were additionally grown in MOPS EZ Rich-glucose media with supplementation of 1 mM aldehydes and aldehyde fate tracked via HPLC after 4 h and **F**. 20 h. Data shown is mean of n=3 with error displayed as standard deviation. The star (*) indicates that the alcohol reductive product for **5** was nondetectable using our HPLC method.

In contrast, and as desired, we detected negligible levels of aldehyde reduction and oxidation for these eight aldehydes at 4 h and 20 h after their addition to cultures of the *E. coli* RARE strain (**Figs. 1E/F**). However, due to the greater volatility of these aldehydes relative to their associated alcohols or acids, we observed greater loss of material when using cells of the RARE strain. This is most noticeable when we supply benzaldehyde (**7**) despite our use of non-breathable seals on the 96 well plates used during both culturing and HPLC sampling. To better examine the case of benzaldehyde addition, we tested growth of RARE cells in capped and sealed 14 mL culture tubes containing 5 mL of media. Under this case, we observed much better preservation of the mass balance, but to our surprise the benzaldehyde had fully oxidized to benzoic acid (**Fig. S1**). Overall, these experiments indicated that most aldehydes of interest are generally stable under aerobic culture conditions, though aldehyde oxidation is possible and a notable concern for benzaldehyde when greater efforts are made to limit its exit from the system.

We next evaluated the stability of these aldehydes when incubated with resting whole cells of the RARE strain (**Fig. 2A**). To prepare resting whole cells, we cultured *E. coli* RARE cells aerobically at 37 °C until mid-exponential phase (OD_600_ = 0.5-0.8), 30 °C for 5 additional hours, and then 18 °C overnight for 18h. Following cell culture, we centrifuged the samples and then resuspended them to a working concentration of 50 mg wet cell weight per mL in 200 mM HEPES buffer, pH 7.0. These methods are similar to commonly used whole cell and buffer concentrations for biocatalysis workflows. We then supplied 1 mM aldehyde and monitored stability. Here, we observed highly unexpected results: All of the aldehydes exhibited some degree of oxidation to the corresponding carboxylic acids, which for most aldehydes meant that complete oxidation occurred within 4 h (**Fig. 2B/C**). To our knowledge, aldehyde oxidation has never been reported as a phenomenon observed when using resting *E. coli* whole cells. This is likely because reduction from aldehyde reductase activity in wild-type strains provided a greater source of aldehyde degradation. Our data from resting cell experimentation shows that the native direction of aldehyde carbon flux can be completely reversed through deletions of genes encoding oxidoreductase enzymes, which suggests that it is not governed by more global variables such as the pool sizes of reducing equivalents. Additionally, the difference we see between aerobic growth and resting cell conditions suggests a difference in the expression level or the apparent concentration of native aldehyde oxidases under different conditions.

**Figure 2.**
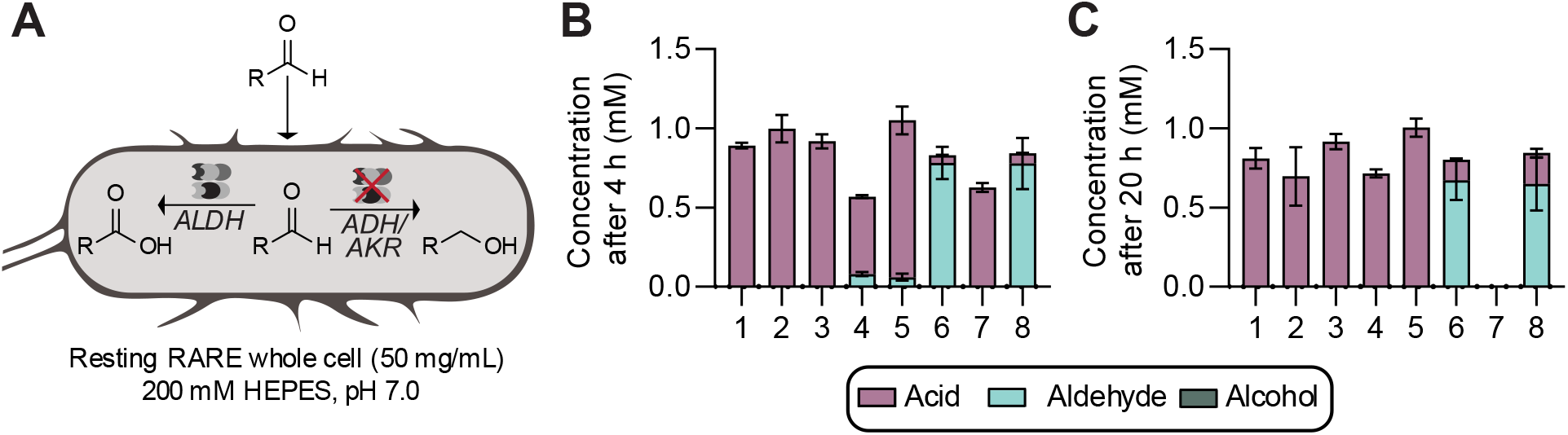
Evaluation of the stability of aldehydes in resting whole cell cultures of *E. coli* RARE. **A**. *E. coli* RARE whole cells were cultured in LB media at 37 °C until mid-exponential phase (OD_600_ = 0.5-0.8), 30 °C for 5 additional hours, and then 18 °C overnight for 18h. Then the cells were centrifuged, washed 2x and resuspended in 200 mM HEPES, pH 7.0 at 50 mg/mL wet cell weight. To initiate experiments, the resuspended whole wells were then supplemented with 1 mM aldehydes. **B**. The stability of aldehydes in resting cells was tracked over 4 h and **C** 20 h. Data shown is mean of n=3 with error displayed as standard deviation.

### 3.2. Construction of a new strain by aldehyde dehydrogenase inactivation decreases oxidation

Given the effectiveness of combinatorial gene deletions at abolishing the reduction of aldehydes and the advancement of multiplexed genome engineering strategies over the last decade, we next investigated whether we could mitigate the observed oxidation using genome engineering. By performing combinatorial gene inactivation using multiplexed automatable genome engineering (MAGE), we could discover the potential identity of the genes responsible for oxidation of these aldehydes and discern whether there are major tradeoffs to inactivating these genes (e.g., fitness or protein expression level). To identify aldehyde oxidase enzymes that could be acting upon our aromatic aldehydes of interest, we performed a BLAST search of the *E. coli* K-12 MG1655 genome for the aldehyde dehydrogenase (ALDH) gene *aldB*, which has previously reported oxidative activity upon aromatic aldehydes. From this search, we identified eleven genes exhibiting similarity to *aldB*, with several reported to be active upon aromatic aldehydes (**Table 1**). We aimed to eliminate the production of AldB and gene products of the five most similar ALDH genes (E-value < 10^−80^): *puuC, betB, patD, feaB*, and *gabD*. We used 10 total rounds of MAGE to inactivate each gene in the *E. coli* RARE strain via introduction of in-frame stop codons within the first 200 codons. We used multiplex allele-specific colony-PCR and confirmation by Sanger sequencing to obtain a variant that contained all six translational knockouts of these ALDHs (**Fig. 3A**).

**Table 1.** Native *E. coli* aldehyde dehydrogenase enzymes identified via BLAST search of *aldB*. All genes shown in bold were investigated for knockout.

| Gene | E-value<br>(*aldB*-basis) | ROAR | Reported activity on aromatic aldehydes |
| --- | --- | --- | --- |
| *aldB* | 0 | ✓ | Benzaldehyde (Ho and Weiner, 2005) |
| ***puuC*** | 5 x 10<sup>-109</sup> | ✓ | Benzaldehyde, Furfural (Jo et al., 2008) |
| ***betB*** | 1 x 10<sup>-101</sup> | ✓ | Benzaldehyde, phenylacetaldehyde (Falkenberg and Strøm, 1990) |
| ***patD*** | 3 x 10<sup>-94</sup> | ✓ | Benzaldehyde, phenylacetaldehyde, *para*-nitrobenzaldehyde (Gruez et al., 2004) |
| ***feaB*** | 9 x 10<sup>-89</sup> | ✓ | Benzaldehyde, phenylacetaldehyde (Rodríguez-Zavala et al., 2006) |
| ***gabD*** | 7 x 10<sup>-81</sup> | ✓ | Benzaldehyde, *meta*-nitrobenzaldehyde, *meta*-carboxybenzaldehyde, *para*-carboxybenzaldehyde (Jaeger et al., 2008) |
| *aldA* | 3 x 10<sup>-79</sup> |  | Benzaldehyde, phenylacetaldehyde (Rodríguez-Zavala et al., 2006) |
| ***sad*** | 9 x 10<sup>-50</sup> |  |  |
| ***astD*** | 9 x 10<sup>-38</sup> |  |  |
| ***putA*** | 3 x 10<sup>-37</sup> |  |  |
| *maoC* | 2 x 10<sup>-16</sup> |  |  |

**Figure 3.**
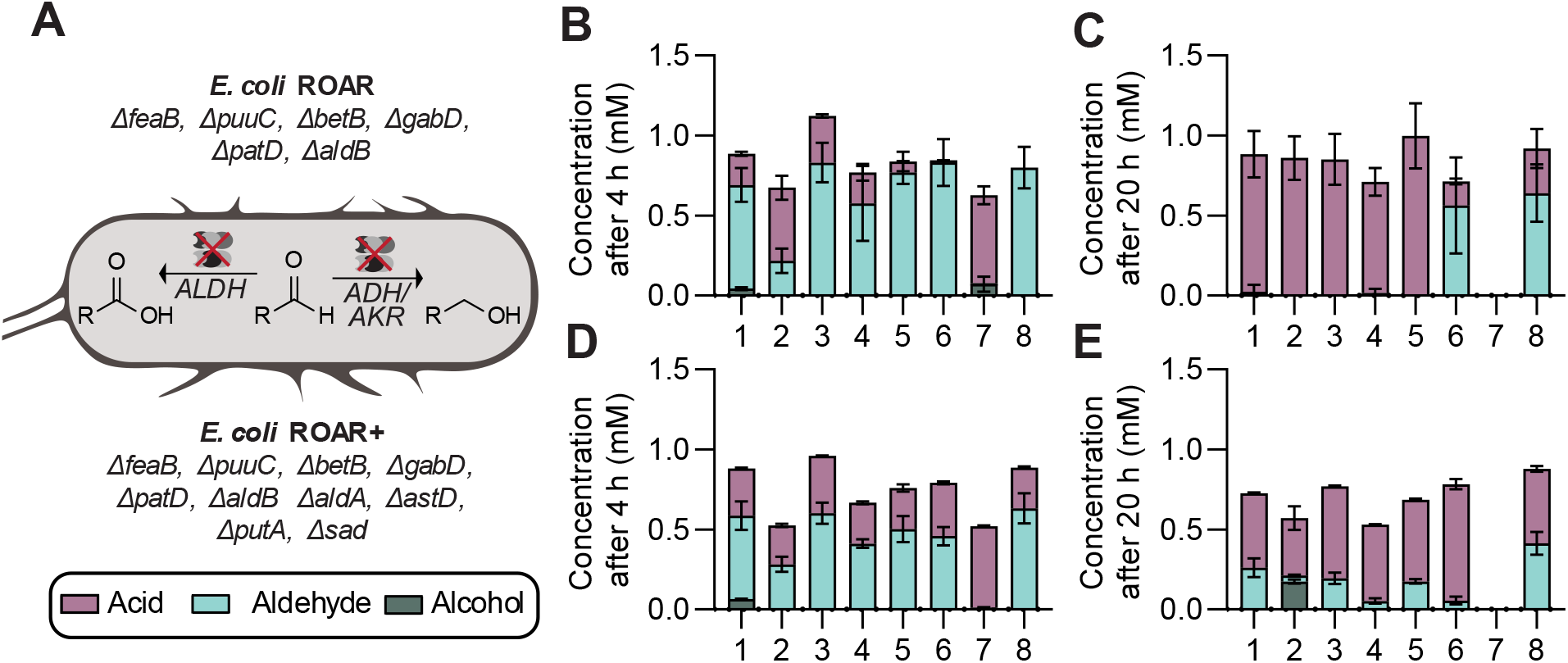
Genomic knockout of aldehyde dehydrogenases (ALDHs) toward improved stability in resting whole cell cultures. **A**. Using the *E. coli* RARE strain as a basis, multiplexed automated genome engineering was performed to translationally knockout six ALDHs (*feaB, puuC, betB, gabD, patD, aldB*) to create a strain for **R**educed **O**xidation **a**nd **R**eduction of aldehydes (ROAR) and a strain with four additional knockouts (*aldA, astD, putA, sad*). **B**. *E. coli* ROAR whole cells were cultured in LB media at at 37 °C until mid-exponential phase (OD_600_ = 0.5-0.8), 30 °C for 5 additional hours, and then 18 °C overnight for 18h. Then, the cells were centrifuged, washed 2x and resuspended in 200 mM HEPES, pH 7.0 at 50 mg/mL. To initiate experiments, the resuspended cells were supplemented with 1 mM aldehydes and the stability of aldehydes in resting cells was tracked over 4 h and **C** 20 h. **D**. *E. coli* ROAR+ resting whole cells supplemented with 1 mM aldehydes and the stability of aldehydes in resting cells was tracked over 4 h and **E**. 20 h. Data shown is mean of n=3 with error displayed as standard deviation.

We set out to determine the stability of the set of aromatic aldehydes in resting whole cells of this new strain. Encouragingly, we noticed that aldehyde oxidation after 4 h decreased for all tested aldehydes (**Fig. 3B**). For a few tested aldehydes, namely piperonal (**1**), *para****-***anisaldehyde (**3**), *meta*-hydroxybenzaldehyde (**4**), and furfural (**5**), the difference in stability using the resting cell preparations of the newly engineered strain rather than the RARE strain was quite stark, with retention of over 50% of the supplied aldehyde after 4 h. When having used RARE resting cells, all aldehydes other than vanillin (**6**) and syringaldehyde (**8**) had depleted to less than 10% of their original value, with several aldehydes undetectable. Given these improvements in aldehyde stability, we named our strain the *E. coli* “ROAR” strain for its <u>R</u>educed <u>O</u>xidation <u>A</u>nd <u>R</u>eduction of aromatic aldehydes. Note that in this study we chose to focus on aromatic aldehydes as they are derived from lignocellulose and amenable to facile detection via HPLC-UV; however, it is possible that inactivation of these promiscuous aldehyde dehydrogenases may offer improvements in stabilization of other kinds of aldehydes as well.

### 3.3. Evaluation of additional ALDH knockouts for possible improved aldehyde stability

For applications that feature resting whole cell catalysis, the time frame of 4 h may be sufficient to achieve complete conversion of catalytic processes, but in some cases, longer reaction times may be necessary to achieve desired conversion. At our later time point of 20 h, unfortunately we observe that resting cells of the ROAR and RARE strains perform comparably poor at stabilizing tested aldehydes (**Fig. 3C**). Thus, we then investigated whether additional aldehyde oxidase knockouts could be beneficial for further improving stability. We selected four additional ALDH genes from our original BLAST search having moderately high similarity to that of *aldB* (E-value < 10^−20^ in this case): *aldA, sad, astD*, and *putA* (**Table 1**). We performed additional cyclic rounds of MAGE to translationally knockout these additional genes. We then evaluated the stability of our aldehyde set in resting whole cells prepared from this strain (ROAR^+^) rather than the ROAR strain at 4 h (**Fig. 3D**) and 20 h (**Fig. 3E**). After 4 h, the ROAR^+^ strain does not appear to provide any stability improvement over the ROAR strain, with potentially greater oxidation observed in some cases. Similarly, after 20 h, retention of aldehydes is quite poor, with greater instability observed in the case of vanillin (**6**) and syringaldehyde (**8**). For three of the aldehydes, piperonal (**1**), *para****-***anisaldehyde (**3**), and furfural (**5**), we do appear to observe some retention after 20 h when using cells prepared from the ROAR^+^ strain. However, because the retention is still quite poor (<25% in all three cases) and because of greater oxidation observed in some cases, it is unclear if these additional knockouts are beneficial overall. Thus, we proceeded ahead with the progenitor ROAR strain instead of the ROAR^+^ strain for further characterization. The superior stabilization afforded by the ROAR strain relative to the RARE strain as measured at 4 h after aldehyde supplementation to resting cells is summarized in **Table 2**.

**Table 2.** Percent of aromatic aldehyde in resting whole cell conditions after 4 h.

| **Aldehyde** | **RARE**<br>**(%)** | **ROAR**<br>**(%)** | **ROAR**<br>***ΔaldA, Δsad,***<br>***ΔastD, ΔputA***<br>**(%)** |
| --- | --- | --- | --- |
| 1 | 0 | 65 ± 11 | 52 ± 9 |
| 2 | 0 | 22 ± 8 | 28 ± 5 |
| 3 | 0 | 83 ± 12 | 60 ± 7 |
| 4 | 8 ± 1 | 58 ± 23 | 41 ± 3 |
| 5 | 6 ± 2 | 77 ± 7 | 50 ± 8 |
| 6 | 78 ± 10 | 83 ± 15 | 46 ± 6 |
| 7 | 0 | 0.50 ± 0.05 | 1.1 ± 0.2 |
| 8 | 78 ± 16 | 80 ± 13 | 63 ± 9 |

While aldehydes had been generally stable under aerobic culturing conditions of the RARE strain, we sought to evaluate whether this stability was maintained in the ROAR strain. Here, we found that in 96 well plate aerobic culturing, aldehyde stability was non-distinguishable between the RARE and ROAR strains (**Fig. S2A-C**). Given that we had previously observed better retention of material and a substantial amount of oxidation when supplying benzaldehyde to cultures grown in culture tubes, we again performed a separate aerobic culturing study in culture tubes (**Fig. S2D-E**). Here, we saw an improvement in benzaldehyde stability when supplying it to cultures of the ROAR strain, with preservation of about 80% of the aldehyde after 20 h compared to the near complete oxidation that had been observed under these conditions when supplying benzaldehyde to cultures of the RARE strain. This result indicates that ROAR may be a preferred host strain for certain applications of aerobic cultures for metabolic engineering or biocatalysis.

Given the improvement in stability that we observed between aerobic culturing conditions and resting cell conditions, we were also curious about aldehyde stability in concentrated cell-free lysate conditions and whether our gene inactivation might lead to improvements there. When harvesting either RARE or RARE cells for preparation of concentrated lysates, we observed a very high level of oxidation within 4 h (**Fig. S3**). This interesting finding indicates that additional genetic modifications may be necessary to inhibit aldehyde oxidation in concentrated lysate, and that until then, cell-free lysate may not be as attractive a platform as resting cells for aldehyde biosynthesis.

### 3.4. Characterization of the ROAR strain for biocatalysis applications

In addition to the metric of aldehyde stabilization, a host strain used for whole cell biocatalysis must exhibit a robust growth rate and an ability to overproduce catalytic proteins of interest. Thus, we investigated whether the ROAR strain could achieve comparable growth rates and protein titers as the RARE strain or its progenitor, the wild-type MG1655 strain. Encouragingly, in complex media such as LB, which is commonly used for protein overexpression, we do not see significant differences in the growth curve or doubling time between the wild-type MG1655 or engineered RARE or ROAR strains (**Fig. 4A**), with doubling times of 20-25 min at 37 °C. In defined media such as M9-glucose minimal media or MOPS EZ Rich-glucose, which can also be optimal for maximal control of media conditions or for growth in industrial scale bioreactors, ROAR appears to grow marginally slower with a doubling time of ∼66 min compared to ∼49 min in MG1655 in M9-glucose and ∼26 min compared to ∼20 min in MOPS EZ Rich-glucose (**Fig. 4B, Fig. S4A**). Next, we investigated the level of production of a superfolder green fluorescent protein (sfGFP) reporter from a plasmid expression vector transformed separately into each of the three strains. We observe comparable OD_600_-normalized production of sfGFP in LB media across all strains tested (**Fig. 4C**). In M9-glucose and MOPS EZ-glycose we observe marginally lower fluorescence arising from the use of ROAR compared to MG1655 (**Fig. 4D, Fig. S4B**). In general, and in resting cell complementary media conditions, ROAR grows and produces protein at levels likely sufficient for biocatalysis compared to MG1655 (doubling times are within 35% for all media tested).

**Figure 4.**
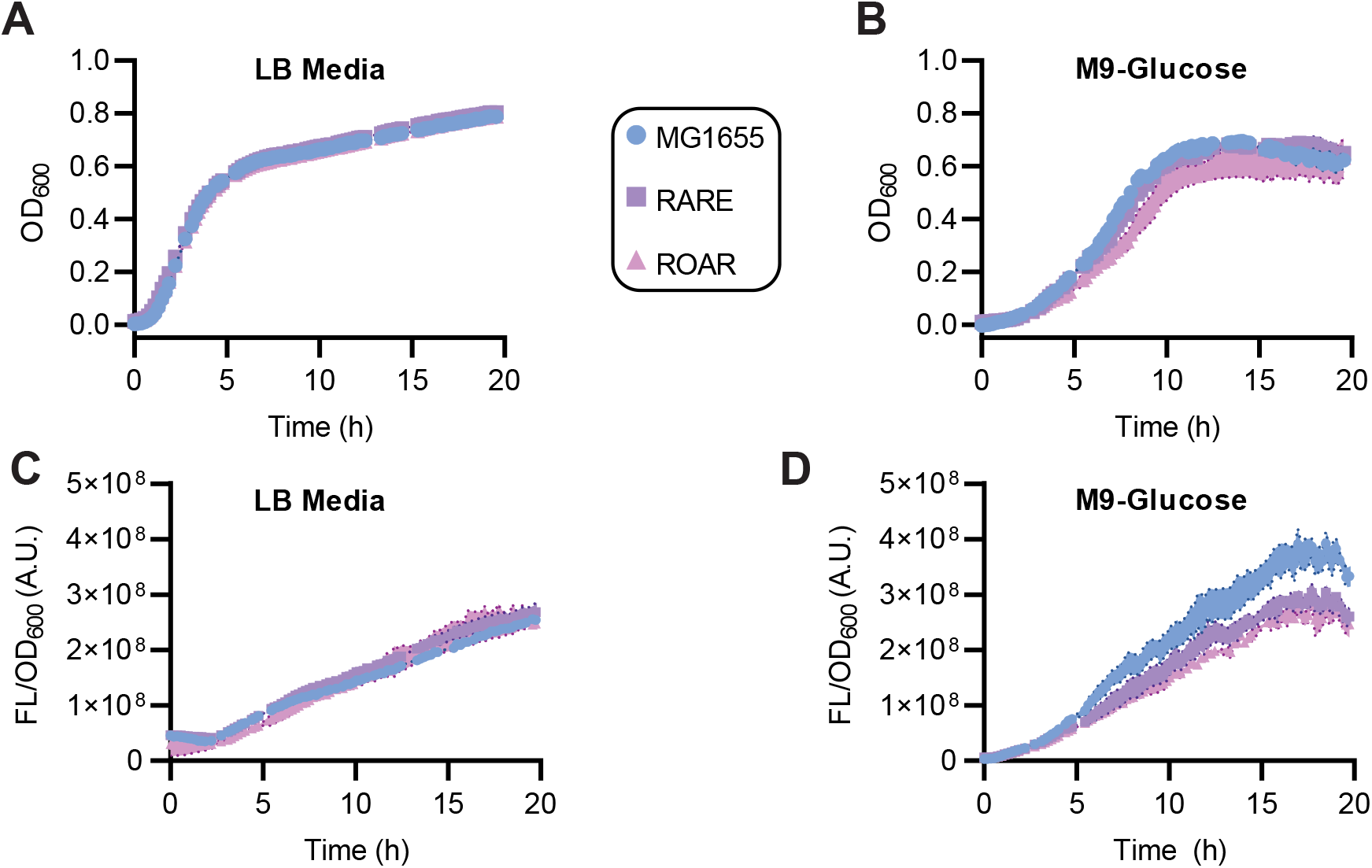
Growth and protein production performance of engineered aldehyde retaining strains. **A**. Growth was monitored via optical density at 600 nm (OD_600_) measured in 96-well plate 20h in LB media and **B**. in M9-glucose minimal media. **C**. Plasmid-based protein overexpression of superfolder green fluorescent protein (sfGFP) was monitored via 96 well plate in a plate reader for 20h by measuring fluorescence (ex: 488 nm, em: 525 nm) normalized by OD_600_ in both LB media and **D**. M9-glucose minimal media. Data shown is mean of n=3 with error displayed as standard deviation.

### 3.5. Application of the ROAR strain for improved aldehyde synthesis

It is anticipated that resting cell preparations of the ROAR strain could be useful for diverse aldehyde biosynthetic processes, and to shed some light on this we examined the biosynthesis of furfural (**5**) from its associated carboxylic acid, furoic acid. These are both versatile platform compounds derived from lignocellulosic biomass, though furfural strongly inhibits bacterial growth. Here, we aimed to use niCAR, an NADPH and ATP-dependent enzyme from *N. iowensis* which has been demonstrated to act upon an array of aromatic aldehydes to perform this bioconversion (**Fig 5A**). To start, we investigated whether niCAR was active upon furoic acid by using an *in vitro* assay. We tracked activity via consumption of NADPH (quantified via measurement of absorbance at 340 nm). Our experiment indicated that niCAR was active on the substrate of interest (**Fig S5**). With this result, we then proceeded to prepare resting whole cells of both the RARE strain and the ROAR strain after transformation with an appropriate plasmid that co-expresses niCAR and the phosphopantetheinyl transferase *Sfp* from *Bacillus subtilis* which is required for CAR activation. We used culturing conditions that maximized niCAR production during fermentation. After harvesting of cells, we resuspended these cells in HEPES buffer and mixed the resting cells with 5 mM of 2-furoic acid substrate and glucose at concentrations of either 10 mM or 100 mM. Glucose is provided to regenerate the cofactors needed by niCAR (**Fig 5B, Fig S6**). We proceeded to monitor production of furfural over time, where we were excited to see that by 4 h the ROAR strain outperformed the RARE strain by over 2-fold in measured furfural titer when using 100 mM glucose (0.70 ± 0.07 mM using RARE, 1.51 ± 0.10 mM using ROAR) or 10 mM glucose (0.78 ± 0.09 mM using RARE, 1.87 ± 0.04 mM using ROAR). When monitoring the reaction for 20 h, we were excited further still to observe that the ROAR strain was able to maintain its production of furfural (1.45 ± 0.26 mM) while reactions in the RARE strain resulted in significant loss of product (0.16 ± 0.06 mM). Thus, use of the ROAR strain achieved a nearly 9-fold enhancement in final furfural titer when using 100 mM glucose. We saw similar average results when providing 10 mM glucose but with greater standard deviation at the 20 h time point. Thus, we have shown that the genetic engineering performed to create the ROAR strain leads to substantial improvements in the stabilization of aldehydes during resting cell conditions and that this improved stabilization can be beneficial during bioconversions that form aldehydes.

**Figure 5.**
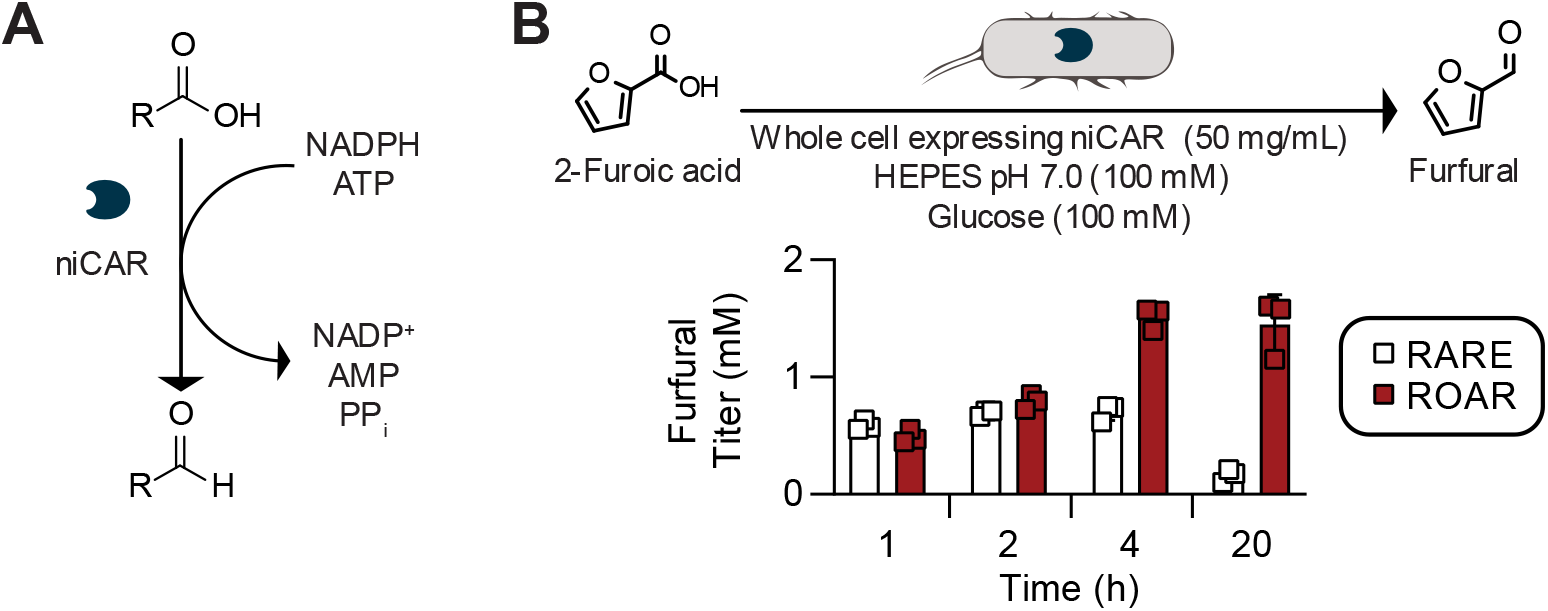
Reduction of furoic acid in resting whole cells expressing *Nocardia iowensis* carboxylic acid reductase (niCAR). **A**. niCAR can reduce aromatic carboxylic acid at the cost of one NADPH and one ATP. **B**. Resting whole cell biocatalysts of RARE or ROAR (50 mg/mL wet cell weight) containing overexpressed niCAR were incubated at 30 °C with 100 mM glucose for cofactor regeneration and 5 mM 2-furoic acid to measure conversion between the two strains. Data shown is mean of n=3 with error displayed as standard deviation.

## 4. Discussion

In this work, we identified that an *E. coli* strain engineered for aldehyde stabilization, the RARE strain, is effective at stabilizing diverse biomass-derived aromatic aldehydes under aerobic growth conditions but does not stabilize aldehydes under resting whole cell conditions. Recently, many researchers have taken to characterizing the activity of aldehyde-associated enzymes, either producers or consumers, in the resting whole cell condition (Doyon et al., 2022; Hepworth et al., 2017). Resting whole cell biocatalysts offer several practical advantages compared to alternatives of fermentation and purified protein biocatalysis, such as simple catalyst separation, no need for cell lysis nor enzyme purification steps, and the ability to tolerate conditions (compounds, solvents, and pH/temperature) that inhibit growth. Our work indicates that previous applications of the RARE strain had not identified aldehyde oxidation because of important differences between aerobic culturing and resting cell conditions, even though our resting cells had been cultured aerobically prior to harvesting. Additionally, unlike in many previous studies, here we did not evaluate aldehyde stability with heterologous expression of highly active CAR enzymes, whose reductive activity can mask the oxidative effects we observed here. For many preferred substrates of CAR enzymes, CAR expression could serve to eliminate perceived oxidation, but this would occur in competition with native aldehyde dehydrogenase activity, leading to futile consumption of ATP and NADPH. An engineering solution to eliminate aldehyde oxidation, as opposed to counteracting it, can avoid this futile cycle.

Here, we used the combinatorial genome engineering technique of MAGE to rapidly terminate the synthesis of full-length ALDHs. Natively, ALDH genes serve several purposes in catabolism and cellular function more broadly with applications in lysine catabolism (glutarate semi-aldehyde dehydrogenase, *gabD*) (Knorr et al., 2018), phenylalanine degradation (phenylacetaldehyde dehydrogenase, *feaB*) (Rodríguez-Zavala et al., 2006), osmolyte synthesis (betaine aldehyde dehydrogenase, *betB*) (Falkenberg and Strøm, 1990), and general toxification of aldehydes (*aldB*) (Ho and Weiner, 2005). Many ALDHs were previously reported to be fairly promiscuous, with activity identified across ranges of both aliphatic substrates and aromatic substrates. For example, AldB is reported to act on aldehydes as distinct as benzaldehyde and acetaldehyde. Given the broad literature-evidenced polyspecificity of these enzymes, we used the speed and ease of MAGE to target different groups of genes, investigating ten different ALDH targets in two different pools of knockouts.

We found that translational knockout of an initial pool of six highly similar ALDHs with reported activity on aromatic aldehydes was sufficient to improve the retention of several aromatic aldehydes as compared to RARE after 4 h. We termed our newly engineered strain that contained the inactivation of all six initial candidate ALDH genes (Δ*aldB*, Δ*puuC*, Δ*betB*, Δ*patD*, Δ*feaB*, and Δ*gabD*) the *E. coli* ROAR strain. We then investigated the knockout of four additional ALDH genes using MAGE, though the improvement in stability afforded by these mutations was minimal, indicating that less obvious aldehyde oxidases could be contributing to stability at longer time scales. Although we did not completely abrogate the phenomenon of aldehyde oxidation, which remains dominant at longer (20 h) time scales, we succeeded in demonstrating substantial improvements in aldehyde stability at shorter (4 h) time scales. These time scales should be more relevant for resting cell biocatalysis processes, especially those that involve aldehyde separation, aldehyde consumption by desired downstream enzymes, or catalyst separation through systems such as flow reactors.

In our previous work to limit the native reduction of aldehydes in metabolically active cells, culminating in the creation of the RARE strain, we knocked out six different aldehyde reducing genes from two distinct families of enzymes, the NADPH-dependent aldo-keto reductase family and the NADH-dependent alcohol dehydrogenase family (Kunjapur et al., 2014). Reduction was significantly decreased for the model aldehydes tested in that work (benzaldehyde and vanillin) and for other aromatic and aliphatic aldehydes tested in follow-on work. Here, we further characterized the RARE strain and noted the stabilization for all eight of the target aromatic aldehydes under aerobic growth conditions. Similarly, in the creation of ROAR, we targeted six genes for knockout which were sufficient to achieve noted improvement in stability, this time limiting oxidation in the resting cell condition. Conversely, in this study, we only targeted gene products from a single family, the NADH-dependent ALDH family of enzymes. Another aldehyde oxidizing enzyme from a different family has been reported in *E. coli* previously, namely the periplasmic, molybdenum-dependent aldehyde oxidase PaoABC, with specificity for aromatic aldehydes (Correia et al., 2016; Neumann et al., 2009). As this enzyme appears to have strict dependence for acidic conditions (pH < 6) in order to exhibit activity, it was not investigated in this work. After creating the ROAR strain, we characterized it for attributes other than aldehyde stabilization and noticed little to no effect on growth or heterologous protein expression, depending on the medium used. As in the case of the RARE strain, this is fortuitous because ALDHs are a highly redundant enzyme class within *E. coli* and are believed to serve important roles in several catabolic pathways. Together, these studies reveal that multiple groups of redundant oxidoreductase enzymes (ADHs, AKRs, and ALDHs) can all be swiftly removed from bacterial cells using combinatorial genome engineering approaches, and that these enzymes are not critical to the cell under a wide range of laboratory conditions. However, unlike in the case of RARE, which afforded nearly complete elimination of aldehyde reduction even at long timescales, the improvements seen in this study are eventually overcome by remaining activity of native enzymes.

Several biosynthetic pathways involving aldehydes as substrates, intermediates, or products can benefit from the improved stability of aromatic aldehydes. Numerous industrially important PLP-dependent enzyme classes including threonine aldolases, threonine transaldolases, and transaminases all utilize aldehydes as substrates for biocatalysis (Doyon et al., 2022; Slabu et al., 2017; Song et al., 2018). Several recent efforts have looked to profile these enzymes in whole cell biocatalysts for ease of catalyst recovery relative to cell-free enzymatic reactions, ease of product separation relative to fermentative contexts, higher throughput screening using cell-based assays on shorter timescales, and greater consistency of production relative to fermentative contexts. In addition, the cost of whole cell catalysts is far lower than purified enzymes and this difference increases as enzyme cascades became longer and require more ancillary enzymes for co-factor or catalyst regeneration.

Here, we demonstrated that the ROAR strain can offer notable improvements for biocatalysis using the example of 2-furoic acid reduction to furfural catalyzed by cells expressing a carboxylic acid reductase. We observed improvements in aldehyde product retention even at long time scales when paired with CAR expression. The reduction of 2-furoic acid to furfural resulted in a two-fold enhancement in furfural titers when using resting cells of the ROAR strain rather than the RARE strain 4 h after reaction initiation and a nearly nine-fold enhancement in furfural titers 20 h after reaction initiation. For an aldehyde end product, extractive protocols can and should be used to help limit loss due to volatility or oxidation and toxicity. However, utilization of a strain with diminished oxidation can provide complementary and dramatic improvements to performance, and we expect this to be more consequential at the higher substrate loadings that would be of greater industrial relevance. Our future work will look to identify whether inefficiencies in biocatalysis using other enzyme classes could be overcome using the ROAR strain.

## Acknowledgements

We acknowledge support from the following funding sources: The National Science Foundation (NSF CMMI-1934887 and CBET-2032243) and minor support from the Center for Plastics Innovation, an Energy Frontier Research Center funded by the U.S. Department of Energy (DOE), Office of Science, Basic Energy Sciences, under Award No. # DE-SC0021166.

## Author Statement

A.M.K. conceived and supervised the study, performed experiments that first created and prototyped the ROAR strain, and helped write the manuscript; N.D.B. designed and conducted all stability experiments, analyzed data, prepared figures, and wrote most of the manuscript; S.R.A. performed the MAGE to create ROAR and performed growth and protein production assays; R.M.D. performed the MAGE to create ROAR+; P.N. initially documented and characterized the phenomenon of aldehyde oxidation in our lab and prepared strains for niCAR testing.

