## Supporting Information for "Combinatorial gene inactivation of aldehyde dehydrogenases mitigates aldehyde oxidation catalyzed by resting cells of *E. coli* RARE strains"

### I. Supplementary Tables

**Table S1.** Strains and plasmids used in this study

| Name | Relevant genotype | Source |
|---|---|---|
| *E. coli* strains | | |
| DH5α | F– Φ80*lac*ZΔM15 Δ(l*ac*ZYA-argF) U169 *rec*A1 *end*A1 *hsd*R17 (rK–, mK+) *pho*A *sup*E44 λ– *thi*-1 *gyr*A96 *rel*A1 | NEB |
| MG1655 | F- λ- *ilv*G- *rfb-50 rph-1* | ATCC 700926 |
| MG1655 (DE3) | F- λ- *ilv*G- *rfb-50 rph-1* (λ DE3)<br>λ DE3 = λ sBamHIo ΔEcoRI-B int::(*lac*I::PlacUV5::T7 gene1) i21 Δ*nin*5 | Previous study (Kunjapur et al., 2014) |
| RARE | MG1655(DE3) Δ*dkgB* Δ*yeaE* Δ*(yqhC-dkgA)* Δ*yahK* Δ*yjgB* | Previous study (Kunjapur et al., 2014) |
| RARE-MAGE | RARE harboring pORTMAGE-Ec1 | This study |
| ROAR | RARE Δ*feaB* Δ*puuC* Δ*betB* Δ*gabD* Δ*patD* Δ*aldB* | This study |
| ROAR+ | ROAR Δ*aldA* Δ*astD* Δ*putA* Δ*sad* | This study |
| NDB01 | MG1655(DE3) harboring pZE-Ub-sfGFP | This study |
| NDB02 | RARE harboring pZE-Ub-sfGFP | This study |
| NDB003 | ROAR harboring pZE-Ub-sfGFP | This study |
| NDB004 | RARE harboring pZE- niCAR-sfp | This study |
| NDB005 | ROAR harboring pZE- niCAR-sfp | This study |
| Plasmids | | |
| pORTMAGE-EC1 | RSF1010 ori, $Kan^R$, harboring CspRecT and *mutL* (E32K) genes | Previous study (Wannier et al., 2020) |
| pZE-Ub-sfGFP | ColE1 ori, $Kan^R$, TetR, Tet promoter with a codon optimized ubiquitin fused to a codon-optimized superfolder GFP | This study |
| pZE-NiCAR-sfp | ColE1 ori, $Kan^R$, TetR, Tet promoter with a codon optimized carboxylic acid reductase gene from *N. iowensis* in an operon with the *sfp* gene from *B. subtilis* | This study |

**Table S2.** Oligonucleotides used in this study. For MAGE oligos, the codon mutations are indicated as (Original codon_Codon position_New codon)

| Oligo Name | Sequence |
|---|---|
| feaB KO oligo (TTA_9_TAA, AGC_10_TGA) | A*A*TAATAAGGAAAAGTGATGACAGAGCCGCATGTAGCAGTATAATGACAGGTCCAACAGTTTCTCGATCGTCAACACGGTCTTTATATTG |
| puuC KO oligo (TAC_8_TAA, TGG_9_TGA) | G*A*CGTGAAACAGGAGTCATAATGAATTTTCATCATCTGGCTTAATGACAGGATAAAGCGTTAAGTCTCGCCATTGAAAACCGCTTATTTA |
| betB KO oligo (GCA_5_TGA, GAA_6_TAA) | T*C*TACCCACCGATTAACCGAGGAGACGTGATGTCCCGAATGTGATAACAGCAGCTTTATATACATGGTGGTTATACCTCCGCCACCAGCG |
| gabD KO oligo (GAA_18_TAA, TGG_19_TGA) | T*T*AACGACAGTAACTTATTCCGCCAGCAGGCGTTGATTAACTGGTAATGACTGGACGCCAACAATGGTGAAGCCATCGACGTCACCAATC |
| patD KO oligo (CAT_3_TAA, TTA_5_TGA) | A*C*AATGGTCAATAACCACTGATACAGGAATATGCTATGCAATAAAAGTGACTGATTAACGGAGAACTGGTTAGCGGCGAAGGGGAAAAAC |
| aldB KO oligo (TTA_21_TGA, AAA_22_TAA) | G*G*GCTACCCATTCGCCGCCAATAAAGTTGTCATAGCGGGCTTATCACTTGAGGGGGAAACCATACTCGCCGGGCTTAATCTGTGCTGAAG |
| putA KO oligo (TGG_194_TGA, TCA_196_TAA) | CAAAATCAGCAACGGTAACTGACAGTAACACATTGGTCGTAGCCCGTCACTGTTTGTTAATGCCGCCACCTGGGGGCTGCTGTTTACTGG |
| aldA KO oligo (GAT_12_TAA, GGA_13_TGA) | A*C*AAACATATAAATCACAGGAGTCGCCCATGTCAGTACCCGTTCAACATCCTATGTATATCTAATGACAGTTTGTTACCTGGCGTGGAGA |
| sad KO oligo (TAT_124_TAA, CGA_125_TGA) | T*C*ACCTGCCATAACGGAAAATTCCACGGCATAATCGCCAGAATCGTCCCCAACGGTCATTACTCAATAACCGCCTGCTGATTTTCCACCA |
| astD KO oligo (TTA_28_TGA, CAA_30_TAA) | G*G*CATCGGCATCATTGCCTTACCATCACACCTCGCCCGATACCGGATTACGCTTCACACGCGATGCGCCCTGGCCCGTTATCCAGTCACC |
| feaB FWD SEQ | CTTTTCTTGTCGCTGCGTACACTG |
| feaB REV SEQ | GTTCCATGGCACAATTCCCGC |
| feaB WT FWD | GACAGAGCCGCATGTAGCAGTATT |
| feaB MUT FWD | GACAGAGCCGCATGTAGCAGTATA |
| feaB REV | GGCAGACATGACTGCGTTATCTACATC |
| puuC FWD SEQ | AGAGGCGGCTTGAAGGATGAAG |
| puuC REV SEQ | GTTCAAATACGCCGCGTGC |
| puuC WT FWD | AGGAGTCATAATGAATTTTCATCATCTGGCTTAC |
| puuC MUT FWD | AGGAGTCATAATGAATTTTCATCATCTGGCTTAA |
| puuC REV | CCAGCTCATGGCTACTGGTGG |
| betB FWD SEQ | TCTGAGCGGCAAACCGCTG |
| betB REV SEQ | CGCGATCGACATCCTCGC |
| betB WT FWD | GGAGACGTGATGTCCCGAATGG |
| betB MUT FWD | GGAGACGTGATGTCCCGAATGT |
| betB REV | GGGCGCTTTTCACGGCG |
| gabD SEQ FWD | CGGCACGTAGTGTGGATGCC |
| gabD WT FWD | GCCAGCAGGCGTTGATTAACG |
| gabD MUT FWD | GCCAGCAGGCGTTGATTAACT |
| gabD REV | GGCACGCTACCCAGCTTGT |
| patD SEQ FWD | GTAACAACCTTGCCGATCCTGGG |
| patD WT FWD | CCACTGATACAGGAATATGCTATGCAAC |
| patD MUT FWD | CCACTGATACAGGAATATGCTATGCAAT |
| patD REV | GGTTTGCCACAATTACGGGACTCCAGT |
| aldB WT FWD | GCCAATAAAGTTGTCATAGCGGGCTTT |
| aldB MUT FWD | GCCAATAAAGTTGTCATAGCGGGCTTA |
| aldB REV | TACAACCTGCCACACGGTGTATG |
| aldB FWD SEQ | TTCAACTATCTCTGTAACCCTTGC |
| aldB SEQ REV | GCAATCTTAAACAGAATCGCCG |
| putA WT FWD | CGGGCTACGACCAATGTGTG |
| putA MUT REV | CGGGCTACGACCAATGTGTT |
| putA REV | CACCATGGGGGTTAAGCTGG |
| putA SEQ FWD | GGAGAGGCTGGCTTCGTTATG |
| aldA WT FWD | AGTACCCGTTCAACATCCTATGTATATCGAT |
| aldA MUT FWD | AGTACCCGTTCAACATCCTATGTATATCTAA |
| aldA REV | AATAGCAGGCAACGCTTCCC |
| aldA SEQ FWD | CGTCTCTCTGATGATTGATGTTAATTAACAATG |
| sad WT FWD | GAATCGTCCCCAACGGTCGA |
| sad MUT FWD | GAATCGTCCCCAACGGTCAT |
| sad REV | GGTATGCAGAACATGGTCCGG |
| sad SEQ FWD | GCGCATGTTTAAGTAAGTAGCCG |
| astD WT FWD | GCATCGGCATCATTGCCTTG |
| astD MUT FWD | GGCATCGGCATCATTGCCTTA |
| astD REV | GGGCAGCCTGCGCAGGG |
| astD SEQ FWD | GGCAAAGCGTTCGACAACGGC |
| pZE backbone FWD | CTTGATGGGGGATCCCATGGTA |
| pZE backbone REV | GTGGTGATGATGGTGATGGCTGCTGCCCATGGTACCTTTCTCCTCTTTAATGAATTCG |
| Car-Sfp FWD | GCCATCACCATCATCACCACATGGCTGTGGACTCGCCG |
| Car-Sfp REV | CCATGGGATCCCCCATCAAGAAGTTACAGCAGTTCTTCGTAGCTCAC |

### **Table S3.** Sequences of proteins expressed in this paper

| Oligo Name | DNA CDS | Protein Sequence |
| --- | --- | --- |
| pZE-Ub-GFP | ATGCAGATTTTTGTGAAGACTTTAACAGGTAAGACGATTACCCTGGAGGTGGAGTCCT<br>CGGACACCATCGATAATGTAAAATCAAAAATCCAAGATAAGGAAGGAATCCCTCCAGA<br>CCAGCAACGTCTGATTTTCGCAGGTAAACAACTGGAGGATGGTCGCACGCTTTCGGAC<br>TACAACATCCAGAAAGAATCTACCCTTCATTTGGTTCTGCGTCTGCGTGGAGGATAGTT<br>GTTTGTGCAGGAGCTTGCATCCAAGGGCGAGGAGCTCTTTACTGGCGTAGTACCAATT<br>CTCGTAGAGCTCGATGGCGATGTAAATGGCCATAAGTTTTCCGTACGCGGCGAGGGC<br>GAGGGCGATGCAACTAACGGCAAGCTCACTCTCAAGTTTATTTGTACTACTGGCAAGC<br>TCCCAGTACCATGGCCAACTCTCGTAACTACTCTGACCTATGGCGTACAATGTTTTTCC<br>CGCTATCCAGATCACATGAAGCAACATGATTTTTTTAAGTCCGCAATGCCAGAGGGCTA<br>TGTACAAGAGCGCACTATTAGCTTTAAGGATGATGGCACCTATAAGACTCGCGCAGAG<br>GTAAAGTTTGAGGGCGATACTCTCGTAAATCGCATTGAGCTCAAGGGCATTGAT**TTT**AA<br>GGAGGATGGCAATATTCTCGGCCATAAGCTGGAGTATAATTTCAATTCCCATAATGTAT<br>ACATTACCGCAGATAAGCAAAAGAATGGCATTAAGGCGAATTTTAAGATTCGCCATAAT<br>GTGGAGGATGGCTCCGTACAACTCGCAGATCATTATCAACAAAATACTCCAATTGGCG<br>ATGGCCCAGTACTCCTCCCAGATAATCATTATCTCTCCACTCAATCCGTGCTCTCCAAA<br>GATCCAAATGAGAAGCGCGATCACATGGTACTCCTGGAGTTTGTAACTGCAGCAGGCA<br>TTACTCATGGCATGGATGAGCTCTATAAGCTCGAGCACCACCACCACCACCACCACTA<br>A | Ub-GFP<br><br>MQIFVKTLTGKTITLEVESSDT<br>IDNVKSKIQDKEGIPPDQQRLI<br>FAGKQLEDGRTLSDYNIQKE<br>STLHLVLRLRGGYLFVQELAS<br>KGEELFTGVVPILVELDGDVN<br>GHKFSVRGEGEGDATNGKLT<br>LKFICTTGKLPVPWPTLVTTLT<br>YGVQCFSRYPDHMKQHDFF<br>KSAMPEGYVQERTISFKDDG<br>TYKTRAEVKFEGDTLVNRIEL<br>KGID**F**KEDGNILGHKLEYNFN<br>SHNVYITADKQKNGIKANFKI<br>RHNVEDGSVQLADHYQQNT<br>PIGDGPVLLPDNHYLSTQSVL<br>SKDPNEKRDHMVLLEFVTAA<br>GITHGMDELYKLEHHHHHHH |
| pZE-niCAR-sfp (entire transcript shown; gene regions are italicized) | GAATTCATTAAAGAGGAGAAAGGTACC*ATGGGCAGCAGCCATCACCATCATCACCACA*<br>*TGGCTGTGGACTCGCCGGATGAACGCCTGCAACGCCGTATCGCCCAACTGTTTGCCG*<br>*AAGATGAACAAGTGAAAGCTGCCCGCCCGCTGGAAGCAGTTAGCGCGGCCGTCTCTG*<br>*CACCGGGTATGCGTCTGGCTCAGATCGCAGCTACGGTGATGGCTGGTTATGCGGATC*<br>*GTCCGGCGGCGGGCCAGCGTGCTTTCGAACTGAATACCGATGACGCAACCGGCCGTA*<br>*CCAGCCTGCGTCTGCTGCCGCGTTTTGAAACCATTACGTACCGCGAACTGTGGCAGC*<br>*GTGTCGGCGAAGTGGCAGCTGCGTGGCATCACGACCCGGAAAACCCGCTGCGTGCG*<br>*GGTGATTTTGTGGCCCTGCTGGGCTTCACCAGCATTGATTATGCAACGCTGGATCTGG*<br>*CTGACATCCATCTGGGTGCGGTTACCGTGCCGCTGCAAGCGAGCGCGGCGGTGTCCC*<br>*AACTGATTGCAATCCTGACCGAAACGAGTCCGCGCCTGCTGGCGTCCACCCCGGAAC*<br>*ATCTGGATGCTGCGGTGGAATGCCTGCTGGCAGGCACCACGCCGGAACGTCTGGTGG*<br>*TTTTCGATTATCACCCGGAAGATGACGATCAGCGCGCCGCATTTGAAAGTGCGCGTCG*<br>*CCGTCTGGCAGATGCAGGTTCCCTGGTGATCGTTGAAACCCTGGACGCGGTGCGTGC*<br>*GCGTGGCCGTGATCTGCCGGCTGCGCCGCTGTTTGTCCCGGATACCGACGATGACCC*<br>*GCTGGCGCTGCTGATTTATACGTCAGGTTCGACCGGCACGCCGAAAGGTGCCATGTA*<br>*CACCAATCGTCTGGCCGCAACGATGTGGCAGGGCAACTCAATGCTGCAAGGCAACAG*<br>*CCAACGCGTTGGCATTAACCTGAATTATATGCCGATGAGTCATATTGCGGGTCGTATCT*<br>*CCCTGTTCGGCGTGCTGGCGCGTGGCGGCACCGCATACTTTGCTGCGAAATCAGACA*<br>*TGAGCACCCTGTTTGAAGATATTGGCCTGGTTCGCCCGACCGAAATCTTTTTCGTTCC*<br>*GCGTGTCTGTGACATGGTGTTTCAGCGCTATCAAAGCGAACTGGATCGCCGTTCTGTC*<br>*GCTGGTGCGGATCTGGACACCCTGGACCGCGAAGTGAAAGCGGATCTGCGTCAGAAT*<br>*TACCTGGGCGGTCGCTTCCTGGTTGCAGTCGTGGGCTCGGCTCCGCTGGCCGCAGAA*<br>*ATGAAAACGTTTATGGAAAGCGTGCTGGACCTGCCGCTGCATGATGGTTATGGCAGTA*<br>*CCGAAGCCGGCGCATCCGTTCTGCTGGATAACCAGATCCAACGTCCGCCGGTCCTGG*<br>*ACTATAAACTGGTCGATGTGCCGGAACTGGGTTACTTTCGCACGGATCGTCCGCACCC*<br>*GCGTGGCGAACTGCTGCTGAAAGCAGAAACCACGATTCCGGGTTATTACAAACGCCC*<br>*GGAAGTTACGGCGGAAATCTTTGATGAAGACGGCTTCTATAAAACCGGCGATATTGTG*<br>*GCCGAACTGGAACATGACCGCCTGGTTTACGTGGATCGTCGTAACAATGTTCTGAAAC*<br>*TGTCCCAGGGCGAATTTGTGACCGTTGCGCACCTGGAAGCTGTGTTCGCGAGCAGCC*<br>*CGCTGATCCGTCAAATTTTTATCTATGGTAGTTCCGAACGCAGTTACCTGCTGGCCGTC*<br>*ATTGTGCCGACCGATGACGCACTGCGTGGCCGCGATACCGCTACGCTGAAAAGCGCT*<br>*CTGGCGGAATCTATTCAGCGTATCGCCAAAGACGCAAATCTGCAACCGTATGAAATTC*<br>*CGCGCGATTTTCTGATCGAAACCGAACCGTTCACGATTGCCAATGGCCTGCTGAGCGG*<br>*TATCGCAAAACTGCTGCGCCCGAACCTGAAAGAACGTTATGGTGCGCAGCTGGAACAA*<br>*ATGTACACCGACCTGGCTACGGGCCAGGCAGATGAACTGCTGGCCCTGCGCCGTGAA*<br>*GCTGCGGATCTGCCGGTGCTGGAAACCGTTAGCCGTGCCGCAAAAGCGATGCTGGGT*<br>*GTGGCAAGCGCGGATATGCGTCCGGACGCACATTTTACCGATCTGGGCGGTGACAGC*<br>*CTGTCTGCACTGAGTTTTTCCAACCTGCTGCACGAAATCTTCGGTGTTGAAGTCCCGG*<br>*TGGGTGTTGTCGTGTCTCCGGCAAACGAACTGCGTGATCTGGCGAATTATATTGAAGC*<br>*CGAACGCAACAGTGGCGCAAAACGTCCGACCTTCACGTCAGTGCATGGCGGTGGCTC*<br>*GGAAATTCGTGCTGCGGATCTGACCCTGGACAAATTTATCGATGCACGCACGCTGGCC*<br>*GCAGCTGATTCTATTCCGCACGCCCCGGTGCCGGCACAGACCGTTCTGCTGACGGGT*<br>*GCGAATGGCTATCTGGGTCGTTTCCTGTGCCTGGAATGGCTGGAACGCCTGGATAAAA*<br>*CCGGCGGCACCCTGATTTGTGTTGTCCGTGGTAGCGACGCGGCGGCGGCACGTAAAC*<br>*GTCTGGATTCAGCCTTTGATAGCGGCGATCCGGGCCTGCTGGAACATTATCAGCAACT*<br>*GGCAGCACGTACCCTGGAAGTGCTGGCAGGCGATATTGGTGACCCGAACCTGGGCCT*<br>*GGATGACGCGACCTGGCAGCGTCTGGCAGAAACGGTCGATCTGATTGTGCATCCGGC*<br>*AGCTCTGGTGAATCACGTTCTGCCGTACACCCAGCTGTTTGGCCCGAACGTGGTTGGC*<br>*ACCGCGGAAATTGTGCGCCTGGCTATCACCGCGCGTCGTAAACCAGTGACCTATCTGT*<br>*CTACGGTTGGCGTCGCAGATCAGGTTGACCCGGCTGAATACCAAGAAGATAGCGATG*<br>*TGCGTGAAATGTCTGCGGTGCGTGTCGTGCGCGAAAGCTATGCCAACGGTTACGGCA*<br>*ATTCTAAATGGGCTGGTGAAGTGCTGCTGCGCGAAGCGCATGATCTGTGCGGTCTGC*<br>*CGGTGGCAGTTTTTCGTTCAGATATGATTCTGGCACACTCGCGCTATGCTGGTCAGCT*<br>*GAATGTCCAAGATGTGTTCACCCGTCTGATTCTGTCACTGGTTGCTACGGGCATCGCG*<br>*CCGTATTCGTTTTACCGCACCGATGCAGACGGTAACCGTCAGCGCGCCCATTACGATG*<br>*GTCTGCCGGCAGATTTCACCGCGGCGGCGATTACGGCGCTGGGTATCCAGGCCACC*<br>*GAAGGCTTTCGCACGTATGATGTGCTGAATCCGTATGATGACGGTATTAGTCTGGACG* | niCAR<br><br><br>MGSSHHHHHHMAVDSPDER<br>LQRRIAQLFAEDEQVKAARPL<br>EAVSAAVSAPGMRLAQIAATV<br>MAGYADRPAAGQRAFELNTD<br>DATGRTSLRLLPRFETITYRE<br>LWQRVGEVAAAWHHDPENP<br>LRAGDFVALLGFTSIDYATLD<br>LADIHLGAVTVPLQASAAVSQ<br>LIAILTETSPRLLASTPEHLDA<br>AVECLLAGTTPERLVVFDYHP<br>EDDDQRAAFESARRRLADAG<br>SLVIVETLDAVRARGRDLPAA<br>PLFVPDTDDDPLALLIYTSGS<br>TGTPKGAMYTNRLAATMWQ<br>GNSMLQGNSQRVGINLNYMP<br>MSHIAGRISLFGVLARGGTAY<br>FAAKSDMSTLFEDIGLVRPTEI<br>FFVPRVCDMVFQRYQSELDR<br>RSVAGADLDTLDREVKADLR<br>QNYLGGRFLVAVVGSAPLAA<br>EMKTFMESVLDLPLHDGYGS<br>TEAGASVLLDNQIQRPPVLDY<br>KLVDVPELGYFRTDRPHPRG<br>ELLLKAETTIPGYYKRPEVTA<br>EIFDEDGFYKTGDIVAELEHD<br>RLVYVDRRNNVLKLSQGEFV<br>TVAHLEAVFASSPLIRQIFIYG<br>SSERSYLLAVIVPTDDALRGR<br>DTATLKSALAESIQRIAKDANL<br>QPYEIPRDFLIETEPFTIANGL<br>LSGIAKLLRPNLKERYGAQLE<br>QMYTDLATGQADELLALRRE<br>AADLPVLETVSRAAKAMLGV<br>ASADMRPDAHFTDLGGDSLS<br>ALSFSNLLHEIFGVEVPVGVV<br>VSPANELRDLANYIEAERNSG<br>AKRPTFTSVHGGGSEIRAADL<br>TLDKFIDARTLAAADSIPHAPV<br>PAQTVLLTGANGYLGRFLCLE<br>WLERLDKTGGTLICVVRGSD<br>AAAARKRLDSAFDSGDPGLL<br>EHYQQLAARTLEVLAGDIGDP<br>NLGLDDATWQRLAETVDLIVH<br>PAALVNHVLPYTQLFGPNVV<br>GTAEIVRLAITARRKPVTYLST<br>VGVADQVDPAEYQEDSDVR<br>EMSAVRVVRESYANGYGNS<br>KWAGEVLLREAHDLCGLPVA<br>VFRSDMILAHSRYAGQLNVQ<br>DVFTRLILSLVATGIAPYSFYR<br>TDADGNRQRAHYDGLPADFT<br>AAAITALGIQATEGFRTYDVL<br>NPYDDGISLDEFVDWLVESG<br>HPIQRITDYSDWFHRFETAIR |

| | | |
|---|---|---|
| | *AATTTGTTGATTGGCTGGTCGAATCCGGCCATCCGATTCAGCGTATCACGGATTATTCA*<br>*GACTGGTTTCACCGCTTCGAAACCGCCATCCGTGCACTGCCGGAAAAACAGCGTCAA*<br>*GCCAGCGTGCTGCCGCTGCTGGATGCATACCGTAACCCGTGTCCGGCCGTTCGCGGT*<br>*GCAATTCTGCCGGCTAAAGAATTTCAGGCTGCGGTCCAAACCGCGAAAATTGGCCCG*<br>*GAACAGGATATTCCGCACCTGAGTGCCCCGCTGATTGATAAATACGTGTCTGACCTGG*<br>*AACTGCTGCAACTGCTGGGTAGTGGCTCTGGACTGGTGGGTGCCCTGATGCACGTGA*<br>*TGCAGAAGCGCAGCCGCGCCATCCACTCCTCCGACGAAGGGGAGGACCAGGCTGGC*<br>*GATGAAGATGAAGATTGA*GCGGCCGCTAATAAAAGGAGATATACC*ATGAAAATCTATG*<br>*GCATTTACATGGATCGTCCGCTGAGTCAGGAAGAAAACGAACGCTTTATGACCTTCAT*<br>*CAGCCCGGAAAAACGTGAAAAATGCCGTCGCTTTTATCATAAAGAAGATGCACACCGC*<br>*ACGCTGCTGGGCGATGTGCTGGTTCGTAGCGTGATCTCTCGCCAGTATCAGCTGGATA*<br>*AATCTGATATTCGTTTCAGTACCCAGGAATACGGTAAACCGTGTATTCCGGATCTGCCG*<br>*GATGCACATTTTAATATCAGCCACTCTGGCCGCTGGGTTATTGGTGCGTTCGATTCTCA*<br>*GCCGATTGGTATCGATATTGAAAAAACGAAACCGATCAGTCTGGAAATTGCCAAACGTT*<br>*TCTTTAGCAAAACCGAATATTCTGATCTGCTGGCAAAAGATAAAGATGAACAGACGGAT*<br>*TACTTTTACCATCTGTGGAGTATGAAAGAATCTTTTATCAAACAGGAAGGCAAAGGTCT*<br>*GAGCCTGCCGCTGGATAGTTTTAGCGTGCGCCTGCATCAGGATGGCCAGGTTTCTATC*<br>*GAACTGCCGGATTCTCACAGTCCGTGCTATATTAAAACCTACGAAGTTGATCCGGGCT*<br>*ATAAAATGGCCGTTTGTGCGGCCCACCCGGATTTCCCGGAAGATATTACGATGGTGAG*<br>*CTACGAAGAACTGCTGTAA*CTTCTTGATGGGGGATCCCATGGTACGCGTGCTAGA | ALPEKQRQASVLPLLDAYRN<br>PCPAVRGAILPAKEFQAAVQT<br>AKIGPEQDIPHLSAPLIDKYVS<br>DLELLQLLGSGSGLVGALMH<br>VMQKRSRAIHSSDEGEDQAG<br>DEDED<br><br>Sfp<br><br>MKIYGIYMDRPLSQEENERF<br>MTFISPEKREKCRRFYHKED<br>AHRTLLGDVLVRSVISRQYQL<br>DKSDIRFSTQEYGKPCIPDLP<br>DAHFNISHSGRWVIGAFDSQ<br>PIGIDIEKTKPISLEIAKRFFSK<br>TEYSDLLAKDKDEQTDYFYHL<br>WSMKESFIKQEGKGLSLPLD<br>SFSVRLHQDGQVSIELPDSH<br>SPCYIKTYEVDPGYKMAVCA<br>AHPDFPEDITMVSYEELL |

### II. Supplemental Figures

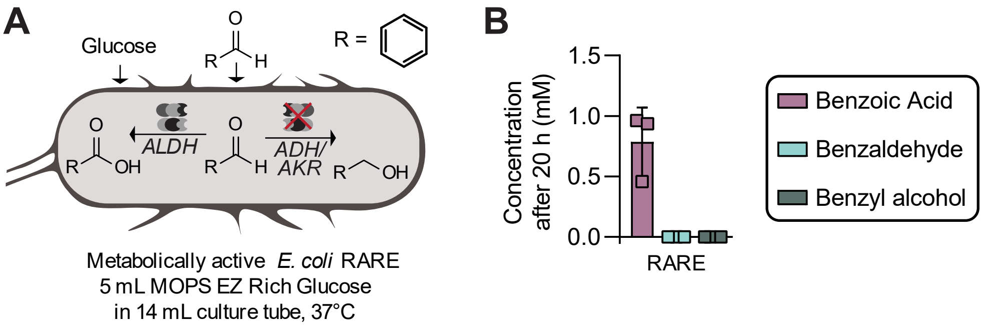


**Figure S1. Evaluation of the stability of benzaldehyde in metabolically active RARE using culture tubes. A.** Benzaldehyde was evaluated for its oxidoreductive stability in metabolically active cultures of *E. coli* RARE using 5 mL of MOPS EZ Rich-glucose media in capped and sealed 14 mL culture tubes at 37 ºC. **B.** Cultures were grown with supplementation of 1 mM benzaldehyde and the fate of benzaldehyde was tracked via HPLC after 20 h. Data shown is mean of n=3 with error displayed as standard deviation

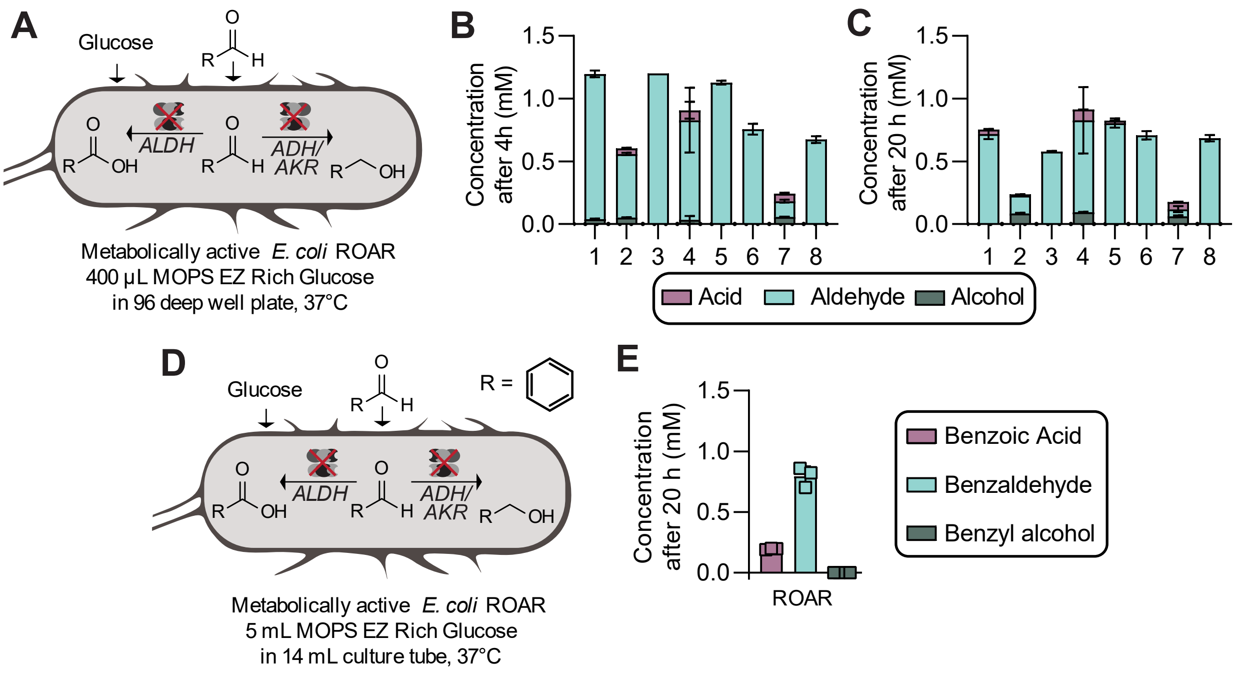


**Figure S2. Evaluation of the stability of aldehydes in metabolically active strains of ROAR. A.** *E. coli* ROAR strains were cultured using 400 µL of MOPS EZ Rich-glucose media in 96 deep well plates at 37 ºC with 1 mM of aldehydes supplemented. **B.** The stability of aldehydes in resting cells in deep well plates was tracked over 4 h and **C** 20 h. **D.** For benzaldehyde, Benzaldehyde metabolically active cultures of *E. coli* ROAR were additionally grown using 5 mL of MOPS EZ Rich-glucose media in 14 mL culture tubes at 37 ºC. **E.** Cultures were grown with supplementation of 1 mM benzaldehyde and the fate of benzaldehyde was tracked via HPLC after 20 h. Data shown is mean of n=3 with error displayed as standard deviation

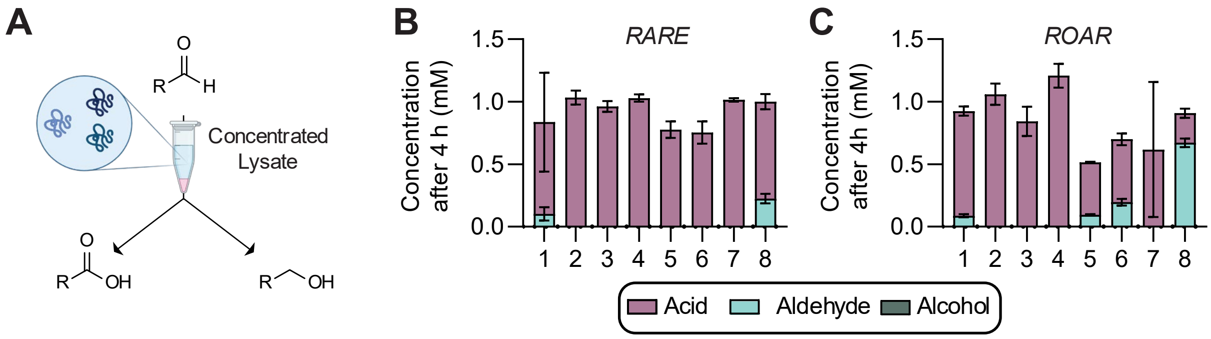


**Figure S3. Evaluation of the stability of aldehydes in concentrated cell-free lysates. A.** *E. coli* cell-free lysates were prepared and concentrated to 5 mg protein/mL in 200 mM HEPES, pH 7.0 at and supplemented with 1 mM aldehydes. **B.** The stability of aldehydes in concentrated cell-free lysate was assayed for RARE and **C.** ROAR. Data shown is mean of n=3 with error displayed as standard deviation

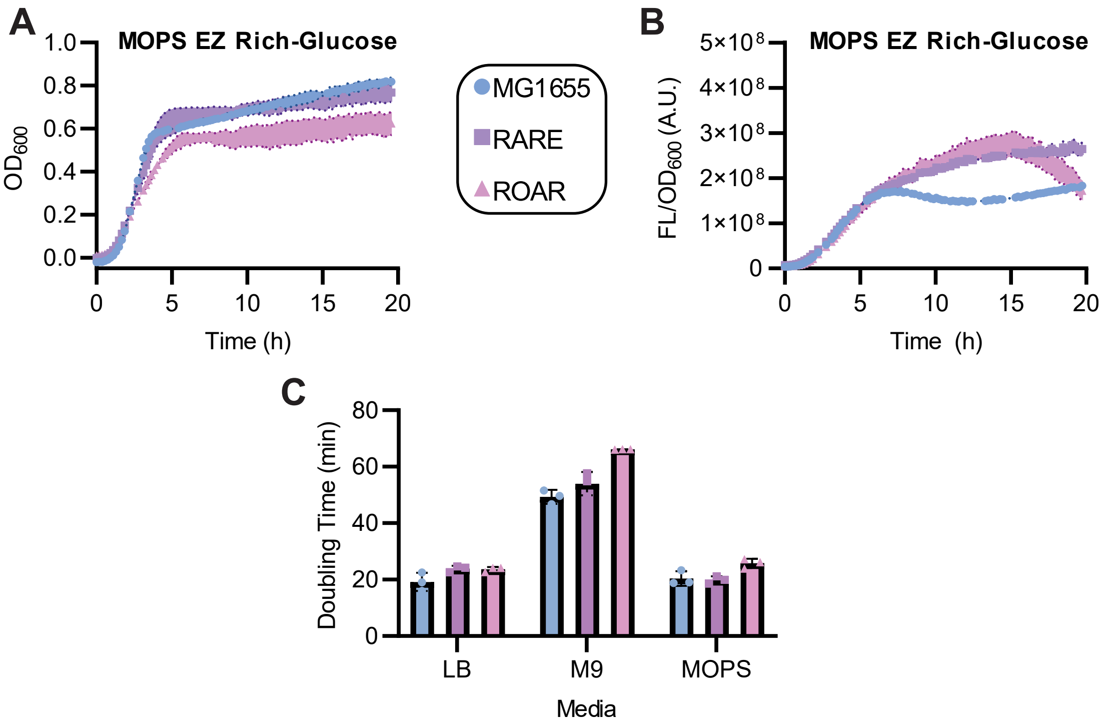


**Figure S4. Growth kinetics in MOPS EZ Rich Media and overall summary. A** Growth was monitored via optical density at 600 nm ($OD_{600}$) measured in 96-well plate 20h in MOPS EZ Rich-Glucose media. **B** Plasmid-based protein overexpression of superfolder green fluorescent protein (sfGFP) was monitored via 96 well plate in a plate reader for 20h by measuring fluorescence (ex: 488 nm, em: 525 nm) normalized by $OD_{600}$ in MOPS EZ Rich-glucose media. **C.** A summary of the maximum doubling times of each strain in each media tested. Data shown is mean of n=3 with error displayed as standard deviation

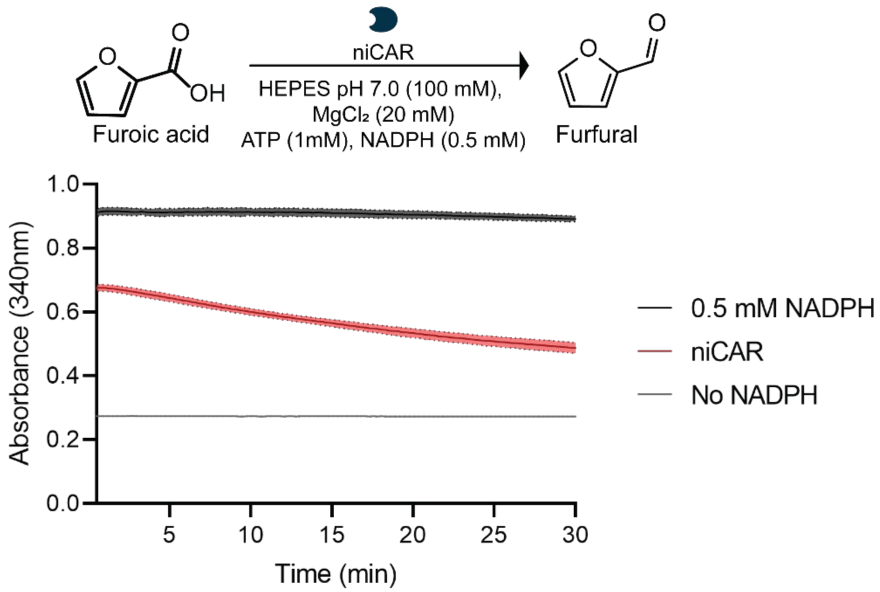


**Figure S5.** ***In vitro* evaluation of niCAR activity on furoic acid.** Carboxylic acid reductase activity was monitored via depletion the absorbance of the cofactor NADPH at 340 nm. Data shown is mean of n=3 with error displayed as standard deviation

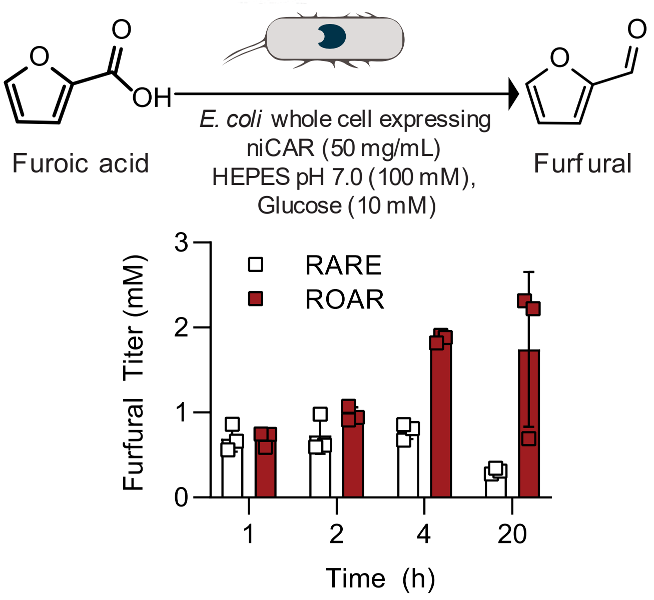


**Figure S6. Reduction of furoic acid in resting whole cells expressing *Nocardia iowensis* carboxylic acid reductase (niCAR).** Resting whole cell biocatalysts of RARE or ROAR (50 mg/mL wet cell weight) containing overexpressed niCAR were incubated at 30 °C with 10 mM glucose for cofactor regeneration and 5 mM 2-furoic acid to measure conversion between the two strains. Data shown is mean of n=3 with error displayed as standard deviation.
